# End-to-end automation of repeat-target cryo-EM structure determination in CryoSPARC

**DOI:** 10.1101/2025.10.17.682689

**Authors:** CryoSPARC Team, Kelly Barber, Hannah Bridges, Suhail Dawood, Katherine Elder, Nick Frasser, Fiona Hu, Serena Liu, Michael McLean, Ryan Narine, Telisha Ng, Valentin Peretroukhin, Ali Punjani, Harris Snyder, Kye Stachowski, Wolfram Tempel, Saara Virani, Rich Waldo, Christina Wang, Nicholas Wong

## Abstract

Single particle cryo-EM is a valuable and growing technique for life science and drug discovery. Currently, obtaining state-of-the-art results from cryo-EM data analysis requires a human in the loop to analyze intermediate results and make image processing decisions. This bottleneck limits the achievable throughput of structure determination, especially in high-throughput settings such as structure-based drug design. In this work, we develop an end-to-end automation strategy for repeat-target structure determination using new tools in CryoSPARC. We demonstrate completely hands-off processing of 21 challenging G protein-coupled receptor (GPCR) datasets. In 17 of 21 cases, automated processing meets or exceeds published resolution and map quality and, in several cases, provides significant improvement in receptor and ligand density that allows improved model building. Our results on both active and inactive state GPCRs show that our automation strategy generalizes easily to new target classes, and that complete automation of data processing is straightforward to achieve in CryoSPARC. We provide downloadable CryoSPARC Workflow files so that users can import, replicate, adapt and extend our automated workflow for their own targets, enabling cryo-EM to be applied at larger scales and to answer larger biological questions.

## 1 Introduction

Single particle cryo-electron microscopy (cryo-EM) has become widely adopted across life science and drug discovery domains globally over the past few years. Cryo-EM sample preparation, data acquisition instruments, image analysis, and model building have all seen substantial improvements in terms of workflow optimization, throughput, and achievable results throughout the last decade. With its applicability to a wide range of targets, cryo-EM is proving to be a highly valuable technique for experimental protein structure determination [1].

As cryo-EM usage expands, scientists seek to tackle more complex biological questions using structural outputs. These questions often require going beyond solving a single structure of a target molecule, to solving multiple structures of a protein complex in different conditions (repeat-target structure determination). Examples include structure-based drug design (SBDD) efforts with multiple ligand-bound structures [2], epitope mapping [3], assessing conformational landscapes through time-resolved studies [4], and validation of structures designed through synthetic biology [5]. SBDD is perhaps the most obvious use case for rapidly determining multiple related structures, where ideally the only variation in sample composition arises from the identity of ligands or binding partners and where the final 3D map resolution and quality directly affect downstream decision-making and the optimization of drug leads.

Currently, the majority of data processing for both single-dataset and repeat-target structure determination is done manually. Various tools exist for streamlining the pre-processing phase of single-particle analysis (SPA) by performing some of these steps on-the-fly or in real time with data collection [6, 7, 8, 9]. Scripting and command-line-based tools are also employed to reduce the number of physical clicks that a scientist needs to perform [10, 11, 12]. However, user-driven analysis and decision making are required at several stages of data analysis in order to obtain state-of-the-art results. Particle picking, particle curation, and final refinements are particularly challenging and often require the most intervention and time. For example, it is typical for users to interactively inspect particle picks, filter based on picking scores, curate training data for neural networks, perform user-guided rounds of classification and selection in 2D or 3D, create masks, and set parameters and search extents for multiple refinement tests.

Having a human in the loop means each data analysis step is done carefully and exactingly, but there are downsides: expert knowledge about cryo-EM and about the target is often required in order to visually analyze and validate intermediate results, to diagnose issues, and to make decisions at each stage. These interventions limit how quickly a final result can be obtained (i.e., time-to-structure), how many structures can be determined in parallel, and also how quickly new scientists can be trained to process and analyze data. Manual processing can therefore become the limiting factor in cryo-EM throughput, despite continuous improvements in instrumentation and data collection speed.

We envision a future where single particle cryo-EM can realize its full potential by becoming routine and scalable, such that it can be employed as the “inner loop” of biological problem solving. This is especially important for SBDD, where the need for cryo-EM at scale is most obviously apparent. A key driver of this evolution will be the automation of single-particle data analysis, to a level that meets or exceeds the quality of manually-processed, human-driven results, even for the most challenging targets.

Development of a method capable of providing this level of automation has been an open challenge in the field, with attempts made by several groups [13, 8, 14, 12, 15, 16, 17]. Most existing approaches center on automatic 2D class selection as a means to curate particles, with results that demonstrate proof-of-concept rather than systematically meeting or exceeding the quality of manual processing. A solution to the challenge of automated processing would need to consistently deliver equal or better results than manual processing, and, for the repeat-target use case, would need to be applicable to real-world datasets, generalize easily to new target classes, and require only limited prior information about the target class.

In this work, we demonstrate a concrete solution for true automation. We describe a generalized automation strategy using a set of existing and newly developed tools in CryoSPARC, which fully automate repeat-target processing. We present automation results on 21 challenging G protein-coupled receptor (GPCR) datasets in both active and inactive states, and for 17 of those datasets we are able to meet or exceed resolution and map quality compared to published results. For the other four datasets, we achieve resolutions within 0.2Å and similar map quality compared to the published depositions. Most importantly, our automated results provide significant improvement in receptor and ligand density for several datasets, allowing for improved modeling of loops, N-termini, and ligands.

All of the requisite jobs and tools for automated processing are available in CryoSPARC v4.7.1. Users can adopt our automation strategy immediately for their own targets, and we provide specific details and instructions in Practical Usage and Automating New Target Classes. The development and availability of fully automated processing will accelerate the pace at which single-particle cryo-EM can scale, and will expedite the transformative impact it can have in life science discovery and medicine.

## 2 GPCRs as a Challenging Test Case for Automation

We demonstrate our automated data processing strategy on G protein-coupled receptors (GPCRs). We chose GPCRs because they represent real-world drug targets that are ubiquitous across both academic and industry research projects, because a significant number of publicly deposited GPCR datasets are available, and because they are notoriously challenging for single-particle analysis.

GPCRs are crucial for transducing extracellular signals such as photons, lipids, metabolites, ions, neurotransmitters, peptides, and proteins to the cytoplasm. This occurs through their interactions with themselves (oligomerization), heterotrimeric G-protein complexes, *β*-arrestins, and GRKs, among others [18]. Given their wide range of ligands and interaction partners, GPCRs are integral to nearly every disease category, from endocrine and metabolic disorders to respiratory and ocular diseases. The human genome contains roughly 800 GPCRs, with approximately 350 non-olfactory GPCRs identified as pharmaceutical targets [19]. As of 2025, approximately 160 GPCRs are the targets for 36% of all FDA-approved drugs [20].

GPCRs (Figure 1) are small (approximately 60 kDa) membrane-bound receptors consisting of seven transmembrane helices and three intracellular and three extracellular loops. Cryo-EM structural characterization of GPCRs has mainly focused on the active state, where a receptor is bound to a heterotrimeric G-protein complex (comprising G*α*, G*β*, and G*γ* subunits) and often times a ligand. In recent years, cryo-EM structural characterization of inactive receptors, typically coupled to a fiducial marker like BRIL or nanobody 6 (Nb6), has become more prevalent. This is due to significant advancements in biochemical approaches and stabilization, data quality, and data processing methods, which have greatly improved access to small, membrane-bound proteins [21, 22].

**Figure 1.**
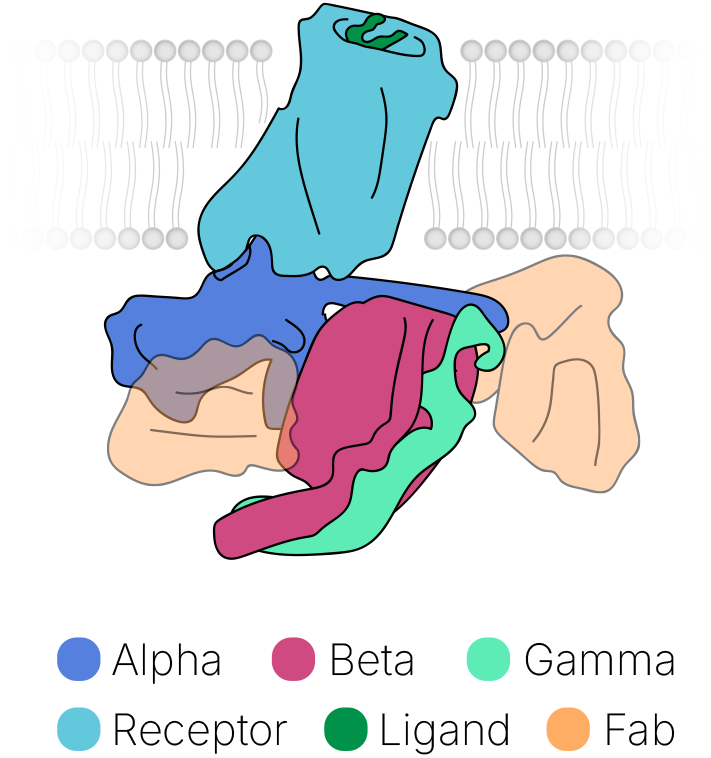
Cartoon diagram of a ligand-bound GPCR embedded in a lipid membrane, showing the receptor, G-protein subunits and ligand. Two possible Fabs (Nb35 and scFv16/30, transparent orange) are shown in their respective locations.

In this work, we apply our automated processing strategy to 21 datasets (Table 1) of GPCR complexes in both the active and inactive state sourced from EMPIAR [23]. Specifically, we used 18 active and 3 inactive state GPCR datasets whose molecular weights range from 61 to 172 kDa. Of the active GPCR datasets, 5 use nanobody 35 (Nb35), 11 use a single chain variable fragment (scFv16 or scFv30) to stabilize the receptor-heterotrimeric G-protein complex, and the last 2 datasets do not have a stabilizing antibody fragment or nanobody present. All active targets have ligands bound: 8 bound to peptidic ligands and the other 10 bound to small molecules. Of the 3 inactive datasets, 2 utilize nanobody 6 (Nb6) as a fiducial marker and one uses megabody 6 (Mb6, derived from Nb6); 2 have small molecule ligands bound while the third is apo. Published resolutions of these targets range from 2.2Å to 3.5Å.

**Table 1.** The 21 EMPIAR datasets used to test our automated processing strategy, categorized into four target classes. The datasets contain a variety of GPCRs and ligands, and they were prepared, collected and originally processed at multiple institutions by multiple teams. Our results demonstrate end-to-end automated processing for all datasets.

| EMPIAR | PDB[24] | EMDB[25] | Target | MW (kDa) | GPCR Class | Ligand Bound |
| --- | --- | --- | --- | --- | --- | --- |
| **Active - Nb35** |  |  |  |  |  |  |
| 10346 [26] | 6ORV | 20179 | GLP-1 | 165 | B | TT-OAD2 |
| 10359 [27] | 6P9Y | 0993 | PAC1-Gs | 167 | B | PACAP-38 (peptide) |
| 10361 [27] | 6P9X | 30003 | CRF1-Gs | 161 | B | CRF (peptide) |
| 10673 [28] | 6X18 | 21992 | GLP-1 Gs | 163 | B | GLP-1 (peptide) |
| 11352 [29] | 7DH5 | 30678 | β3-adrenergic recept. | 133 | A | Mirabegron |
| **Active - scFv16/30** |  |  |  |  |  |  |
| 10288 [30] | 6N4B | 0339 | CB1-Gi | 171 | A | MDMB-Fubinaca |
| 10853 [31] | 7S8L | 24896 | MRGPRX2-Gq | 140 | A | Cortistatin-14 (peptide) |
| 10854 [31] | 7S8N | 24898 | MRGPRX2-Gq | 139 | A | (R)-ZINC-3573 |
| 10855 [31] | 7S8M | 24897 | MRGPRX2-Gi | 172 | A | Cortistatin-14 (peptide) |
| 10856 [31] | 7S8O | 24899 | MRGPRX2-Gi | 170 | A | (R)-ZINC-3573 |
| 10936 [32] | 7RYC | 24733 | OTR-Gq | 153 | A | Oxytocin |
| 11119 [33] | 7WU9 | 32824 | EP3-Gi | 150 | A | Prostaglandin E2 |
| 11350 [34] | 7YU3 | 34097 | LPA1-Gi | 158 | A | ONO-0740556 |
| 11435 [35] | 7RAN | 24378 | 5-HT2AR-mini Gq | 142 | A | (R)-69 |
| 11448 [36] | 7SRR | 25402 | 5-HT2BR-mini Gq | 146 | A | LSD |
| 11474 [37] | 8GHV | 40052 | CB1-Gi | 170 | A | AMG315 |
| **Active - no Fab** |  |  |  |  |  |  |
| 11432 [38] | 7SF7 | 25076 | LPHN3 (ADGRL3) | 106 | B | Tethered agonist peptide |
| 11433 [38] | 7SF8 | 25077 | ADGRG1-miniG13 | 107 | B | Tethered agonist peptide |
| **Inactive** |  |  |  |  |  |  |
| 11134 [39] | 7UL5 | 26592 | SSTR-Nb6 | 61 | A | N/A |
| 11135 [39] | 7UL4 | 26591 | MOR-Mb6 | 104 | A | Alvimopan |
| 11136 [39] | 7UL2 | 26589 | hNTSR1-Nb6 | 62 | A | SR48692 |

To produce test cases for repeat-target automation, we categorized the 21 datasets into four target classes (Active no-Fab, Active Nb35, Active scFv, and Inactive; see Table 1). Within each target class, we enable fully automated processing, with no human intervention or parameter changes required. It is important to note that our test target classes each contain a wide mixture of GPCR species, ligands, sample preparation parameters, microscopes, and data collection parameters. This makes our tests even more challenging than a typical repeat-target scenario in practice, where the protein species and imaging parameters would be held constant and only the ligands would be varied across datasets.

## 3 Development of a Single-Particle Automation Strategy

Successful automation of the repeat-target scenario for a given target class requires a strategy that can start from a large uncurated dataset of raw cryo-EM exposures and produce high-resolution density maps of the target molecule (including any particular local regions of interest). Furthermore, the strategy should be applicable to challenging real-world datasets and should generalize easily to new target classes. When applying the strategy to a new target class, only limited prior information about the target class should be required.

In this repeat-processing context, it is reasonable to assume that at least one, potentially small or low quality, exemplar dataset from the target class has been manually processed previously; this gives a starting point of prior knowledge to work with in the development of an automation strategy. In particular, in this work we assume that a low resolution (15Å) reference map is available that defines the broad shape of the target class of molecules, and also that masks (relative to the reference) are available that define regions of interest (e.g., a receptor containing a ligand binding pocket). As mentioned in the previous section, we categorized our 21 test datasets into four classes based on their state (active or inactive) and the type of Fab bound (Nb35, scFv, no Fab).

Our automated processing strategy (Figure 2) generally follows the current best practices for manual processing. However, when developing the strategy, we found that some stages required special attention or the creation of new methods and tools in order for automated processing to achieve results equal or superior to manual processing. Micrograph pre-processing (i.e., motion correction, CTF estimation, micrograph curation) are well established and highly automated, but in the unattended automation context we needed to be sure that the workflow would not be derailed by large numbers of poor quality, off-target, or contaminated micrographs. When picking particles, our strategy needed to work well across particle sizes and shapes and not miss rare or low-contrast views, especially for small challenging targets such as GPCRs in low signal-to-noise images. Furthermore, picking needed to work well across changing conditions between datasets such as the sample (target and ligands), presence of detergents or additives, variation in electron dose, magnification, etc. One of the most challenging stages was curating initial picks into a smaller, high-quality particle set without introducing orientation bias, as our GPCR test datasets (like most real-world datasets) contained substantial numbers of broken, denatured or dissociated particles mixed in with rare views. Finally, in order for 3D refinements to yield the maximum map quality and resolution, our workflow needed to make use of CTF refinement and reference based motion correction.

**Figure 2.**
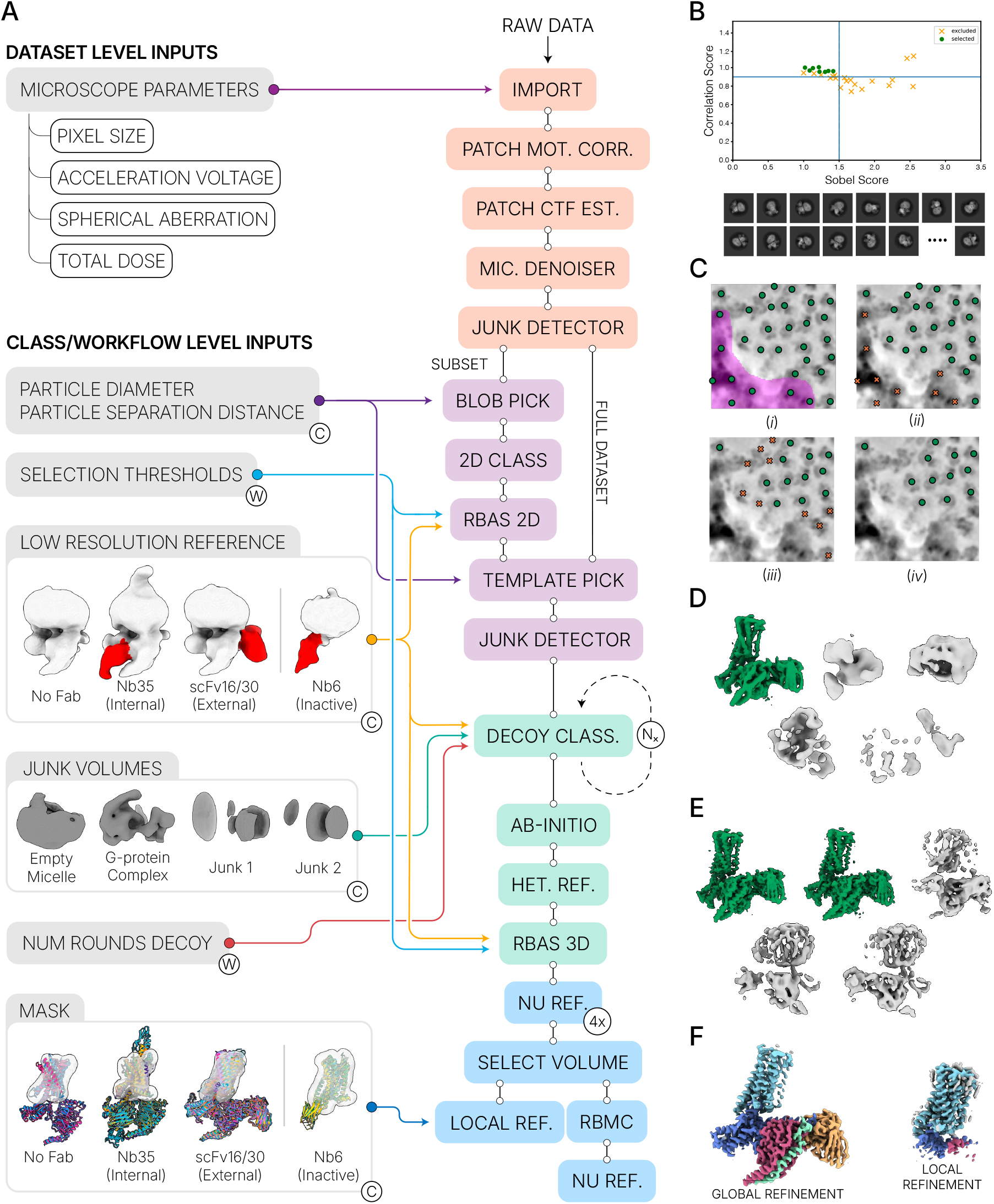
Complete workflow for end-to-end automation of repeat-target structure determination. **A.**Workflow flowchart from import to final refinements, detailing required inputs at each stage. **Dataset level inputs** must be provided for each new dataset. **Class level inputs (C)** only need to be set once, and are reused for all datasets in the target class. For example, the low resolution reference map and corresponding mask for each target class in this work are shown in insets. **Workflow level inputs (W)** can be reused across multiple classes. **B**. 2D classes for template picking are automatically selected using Reference Based Auto Select 2D (RBAS 2D). **C**. Micrograph Denoiser improves particle picking, and particles close to junk and contaminants are automatically rejected by Micrograph Junk Detector. **D**. Decoy classification curates particles in 3D without introducing orientation bias. **E**. Further curation using Ab-initio Reconstruction and Heterogeneous refinement yields final high-quality particles. **F**. Non-Uniform Refinement (left) produces an optimal final global refinement, and Local Refinement (right) using the input mask improves map quality in the receptor region.

The automation strategy we ultimately produced is presented as a flowchart in Figure 2. Each stage of the workflow and each new method or tool is described in detail in Section 7: Methods and Extended Results. At a high level, the workflow pre-processes the data and uses new tools to automatically remove noise and identify contaminants in the micrographs. These steps enable simplified and generalizable particle picking (Figure 2B and Figure 2C). Particle curation is performed entirely in 3D rather than 2D, allowing rare views to be retained while filtering out poor quality particles. The workflow first uses the low-resolution reference and junk volumes to select for high-quality particles even when they are far outnumbered, an approach often referred to as decoy classification (Figure 2D). It ends with ab-initio reconstruction and classification to ensure that dataset-specific signal is used for final curation (Figure 2E). Multiple refinements are then carried out to determine the optimal parameters for CTF refinement and whether or not the dataset will benefit from reference based motion correction. Finally, automatic alignment of the 3D maps enables local refinement to produce maps with maximum quality and resolution in regions of interest such as ligand binding domains (Figure 2F).

To use the automated workflow in practice, there are three categories of inputs that must be provided: dataset level, class level, and workflow level inputs (Figure 2A). Dataset level inputs must be varied with each input dataset; these are microscope parameters, such as the pixel size and electron dose. Target-class level inputs must be determined once at the time of instantiating the automation workflow for a specific target class, and then do not need to be changed; these are the low-resolution (15Å) reference map, mask(s), junk volumes, particle diameter, and minimum particle separation distance. In our experiments, we instantiated the workflow once for each of our four target classes. Finally, workflow level inputs can be held constant across multiple target classes; these are the number of rounds of decoy classification and the selection thresholds for 2D and 3D reference-based auto selection. In our experiments, we held these inputs constant across all active GPCR classes, and changed them for inactive classes (see Section 4.1).

We use the Workflows feature (Figure 3, released in CryoSPARC v4.4) to encapsulate our automated processing strategy, enabling one-click processing from end-to-end. Standard manual processing in CryoSPARC requires a user to interact with the UI to build each job, modify parameters, and connect inputs. The Workflows feature automates this process; a chain or tree of connected jobs (including inputs and parameters, with annotations and notes) can be saved as a Workflow, and re-used on another dataset in a single click. Workflows can be exported in a simple JSON file format and imported into another CryoSPARC instance for easy sharing and replication. For all results shown in this work, we set up a Workflow once for each of the four target classes, and within each class, we did not vary any inputs except the dataset level inputs shown in Figure 2. We have made our Workflow files available to readers to download and re-use; see Section 6.

**Figure 3.**
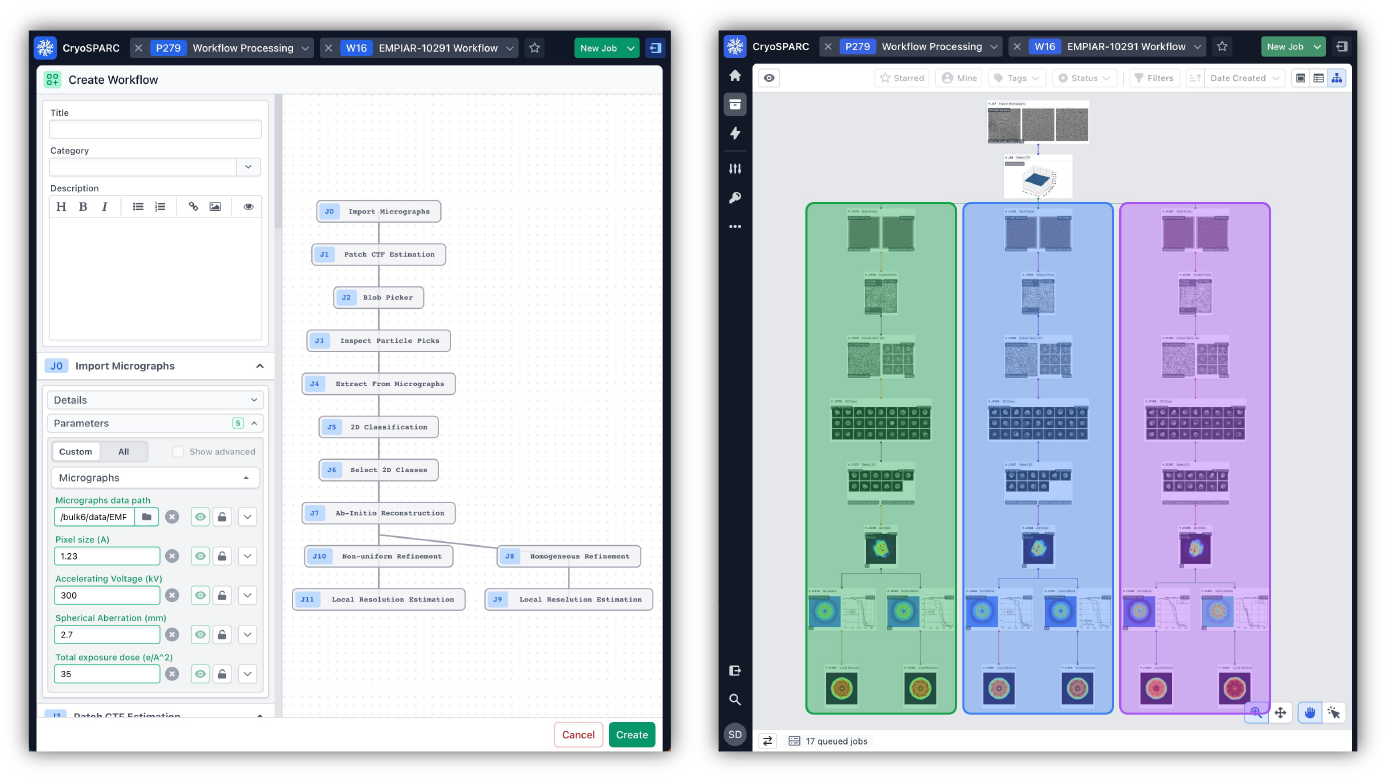
CryoSPARC’s Workflows feature encapsulates our automated processing strategy so it can be executed in one click. A Workflow records a series of connected jobs and can be saved and applied to a new dataset within the UI. Workflows can be exported in JSON format and imported into another CryoSPARC instance for easy sharing and replication. **Left:** Parameters, descriptions and annotations can be saved when creating a Workflow. **Right:** CryoSPARC Tree view showing application of a Workflow multiple times.

Section 7: Methods and Extended Results describes all the stages of our automated workflow in detail, including how we developed each stage and the associated new tools and methods in CryoSPARC: the Micrograph Junk Detector, Micrograph Denoiser, automatic Inspect Particle Picks, Reference Based Auto Select 2D, Reference Based Auto Select 3D, Select Volume, and Align 3D Maps. Section 7 also includes details of the experiments we carried out to determine how variations in the strategy affect processing results. In the next section, we demonstrate the power of our automation strategy across 21 GPCR test datasets.

## 4 Automated Processing Results on 21 GPCR Datasets

In the preceding section and in Methods and Extended Results, we describe the development of a generalizable strategy for complete automation of the processing of repeat-target single-particle cryo-EM datasets. We employed the strategy to automatically process all 21 GPCR datasets described in Section 2. The results, presented in this section, demonstrate that we have successfully reached the goal of being able to produce maps with resolution and map quality equal to or better than manual processing in the repeat-target scenario, with no manual intervention required. We show that the strategy enables unattended processing of datasets containing poor-quality micrographs, contaminants, aberrations, aggregation, denatured, broken and junk particles. Despite the broad variation in input datasets, we are able to automatically curate and isolate sets of good particle images. In multiple cases our fully automated processing results yield improved map quality and interpretability in the receptor region where a ligand is bound, which is a crucial result for drug discovery workflows. The sections and figures below explain how the results were produced and compare them to the originally deposited manual processing results.

### 4.1 Data and Input Preparation

Automated processing requires setting dataset-level, target-class-level and workflow level inputs (Figure 2). Dataset-specific microscope parameters for each dataset were obtained from the respective EMPIAR depositions and related publications, and were entered when importing each dataset into CryoSPARC (Table S1). As mentioned in Section 2, the 21 datasets were categorized into four target classes (Active no-Fab, Active Nb35, Active scFv, and Inactive). Within each target class, all datasets were processed automatically with a fixed set of target-class level inputs (as described in Figure 2). The low-resolution (15Å) reference maps and corresponding masks (shown in Figure 2A) were generated from previous manual processing of a single exemplar dataset for each class (see Supplementary Materials for details). Generating reference maps from a single exemplar means that in practice, for a new target class, an automation workflow can be instantiated after processing only a single dataset manually. Four low resolution junk volumes (shown in Figure 2A) were generated from an exemplar dataset, with one containing an empty micelle, one containing a detached G-protein complex, and two randomly generated densities (see Supplementary Materials for details). The same junk volumes were used for all four target classes. The other target-class level inputs were set as follows: for the three Active classes, blob picking diameter of 110Å (min) to 140Å (max) with separation distance of 0.7 and template picking diameter of 120Å with separation distance of 0.6; for the Inactive class, blob picking diameter of 100Å (min) to 120Å (max) with separation distance of 0.7 and template picking diameter of 110Å with separation distance of 0.6.

Workflow level inputs were held constant across all the Active classes, and changed slightly for the Inactive class. One round of decoy classification was used for Active classes and two for the Inactive class. Selection thresholds for 2D reference-based auto selection remained unchanged across all four classes: correlation 0.9 and Sobel 1.5. The threshold for 3D selection was set to 90% of best resolution for active classes and 95% for inactive classes.

### 4.2 Improved FSC Resolution and Map Anisotropy

Numerical map resolution and isotropy estimates are the simplest way to compare our automated results to deposited manual processing results from the EMDB [25]. Figure 4 displays a comparison of goldstandard FSC (Fourier Shell Correlation) resolution and cFAR (Conical Fourier shell correlation Area Ratio [40]) anisotropy score across the test set, which are also detailed in Table S2. For each dataset, we automatically generated a generous solvent mask that was used to compute FSC resolutions from our half-maps as well as the published deposited half-maps where available, ensuring fair comparison of resolutions (see Supplementary Materials for details). In all 21 test cases, our automation strategy produced density maps of high resolution and quality; in 17 cases, our results matched or improved the resolution over manual processing, in three cases produced results within 0.1Å, and in one case within 0.2Å. In the best six cases, automated processing substantially improved resolution: by 0.7Å, 0.6Å and 0.4Å in EMPIAR-11350, EMPIAR-11435 and EMPIAR-11474 respectively, and by 0.3Å in three additional cases. Furthermore, automated processing improved the cFAR score in 13 of the 15 cases where published half-maps were available for comparison, indicating that automated particle curation typically retained a more isotropic distribution of particle orientations.

**Figure 4.**
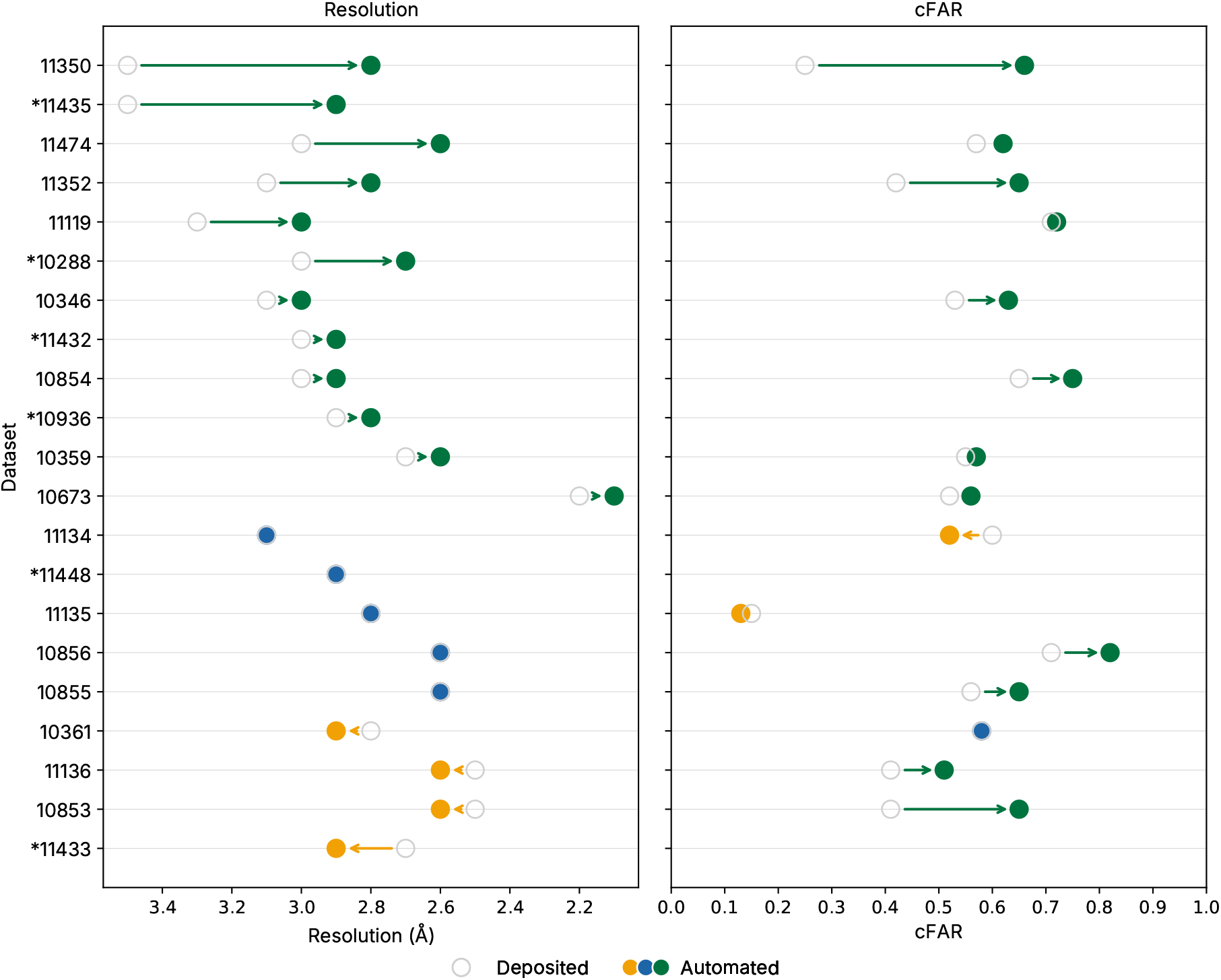
Automated processing matches or exceeds the map resolution and map isotropy compared to deposited manual processing results for majority of GPCR datasets. **Left:** Gold-standard Fourier Shell Correlation (FSC) and **right:** Conical FSC Area Ratio (cFAR). Values from deposited published results shown as gray circles. Values from automated processing shown as filled, colored circles: improved (green), equivalent (blue), or worse (orange). Our automated workflow achieves equal or better resolution in 17 out of the 21 datasets with six improvements of 0.3Å or greater. FSC and cFAR were measured using a single soft mask for each dataset, so that the values are directly comparable (see Section 8.4). Asterisks indicate datasets for which half-maps were not deposited to EMDB, so the deposition-reported resolution is used instead and cFAR scores could not be calculated.

Figure 5 displays, for each test dataset, the global and local density maps resulting from automated processing, as well as the originally published manually processed maps for comparison. Sharpened maps are displayed, with B-factors chosen in each case to improve map quality but retain connectivity. Figure 5 also displays the ligand density visible in our automatically processed maps and in the published maps, for comparison of local map quality. In all cases, local refinement in our automated workflow improved map quality in the receptor domain and produced clear, interpretable ligand density.

**Figure 5.**
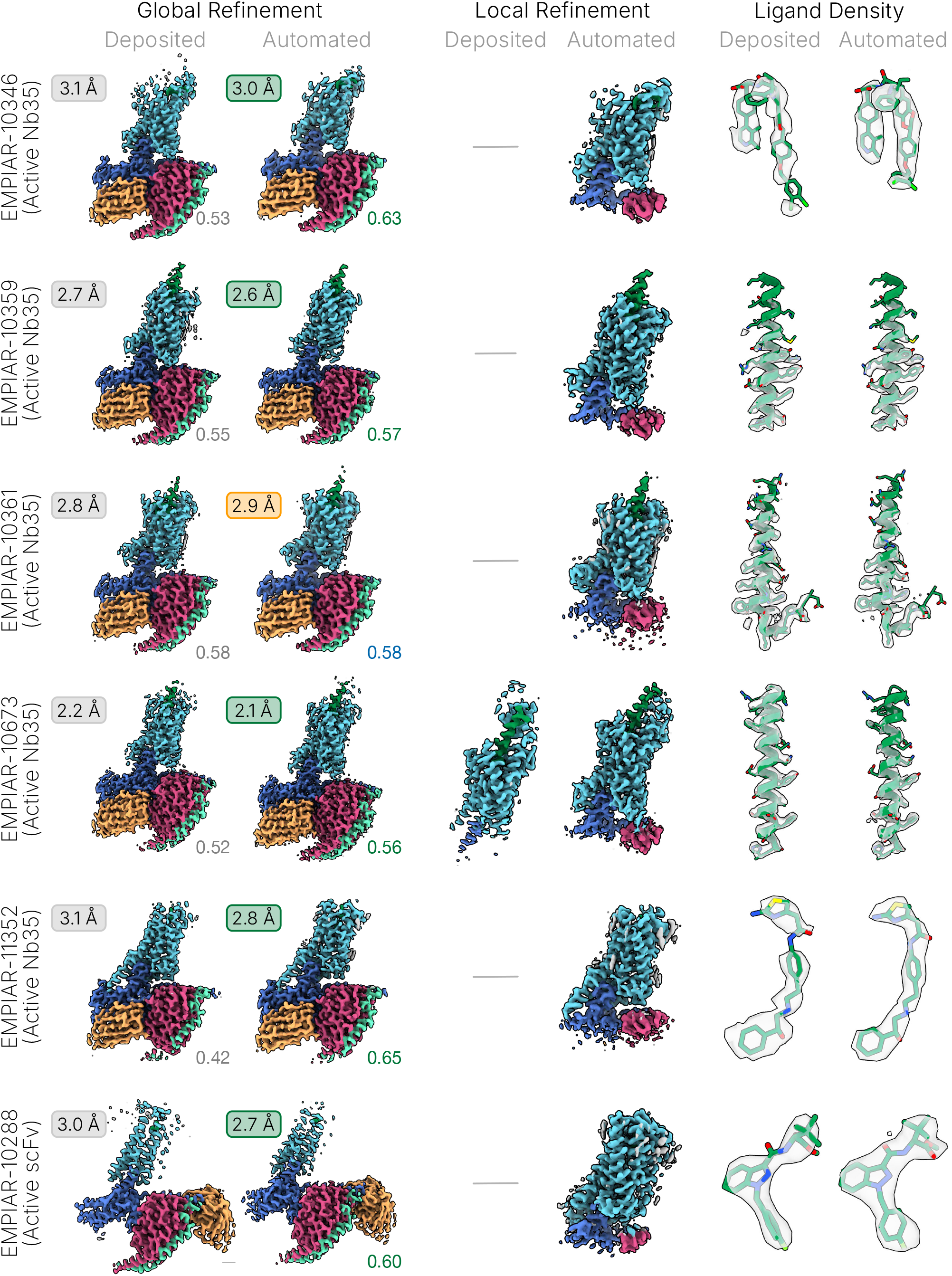
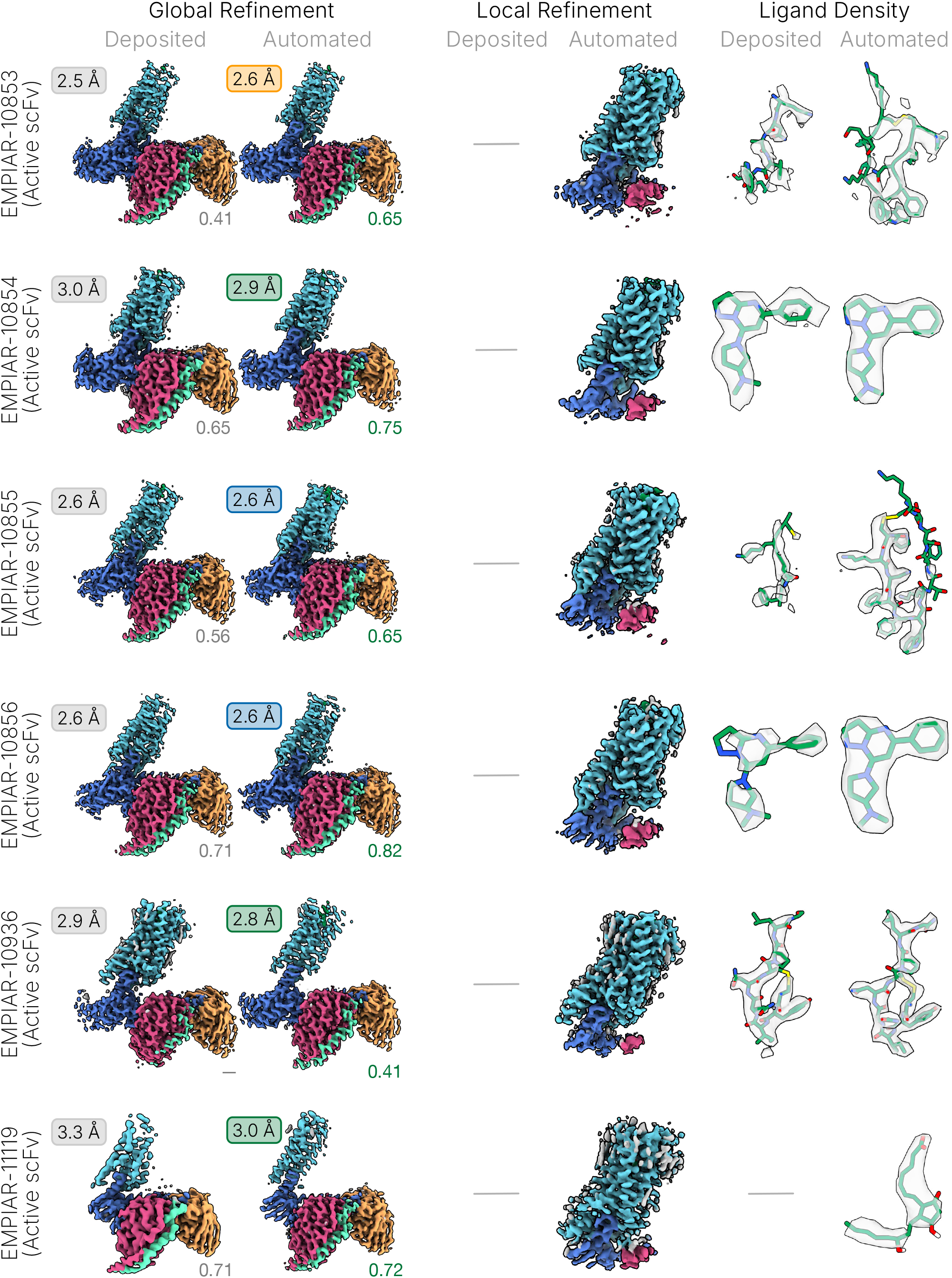
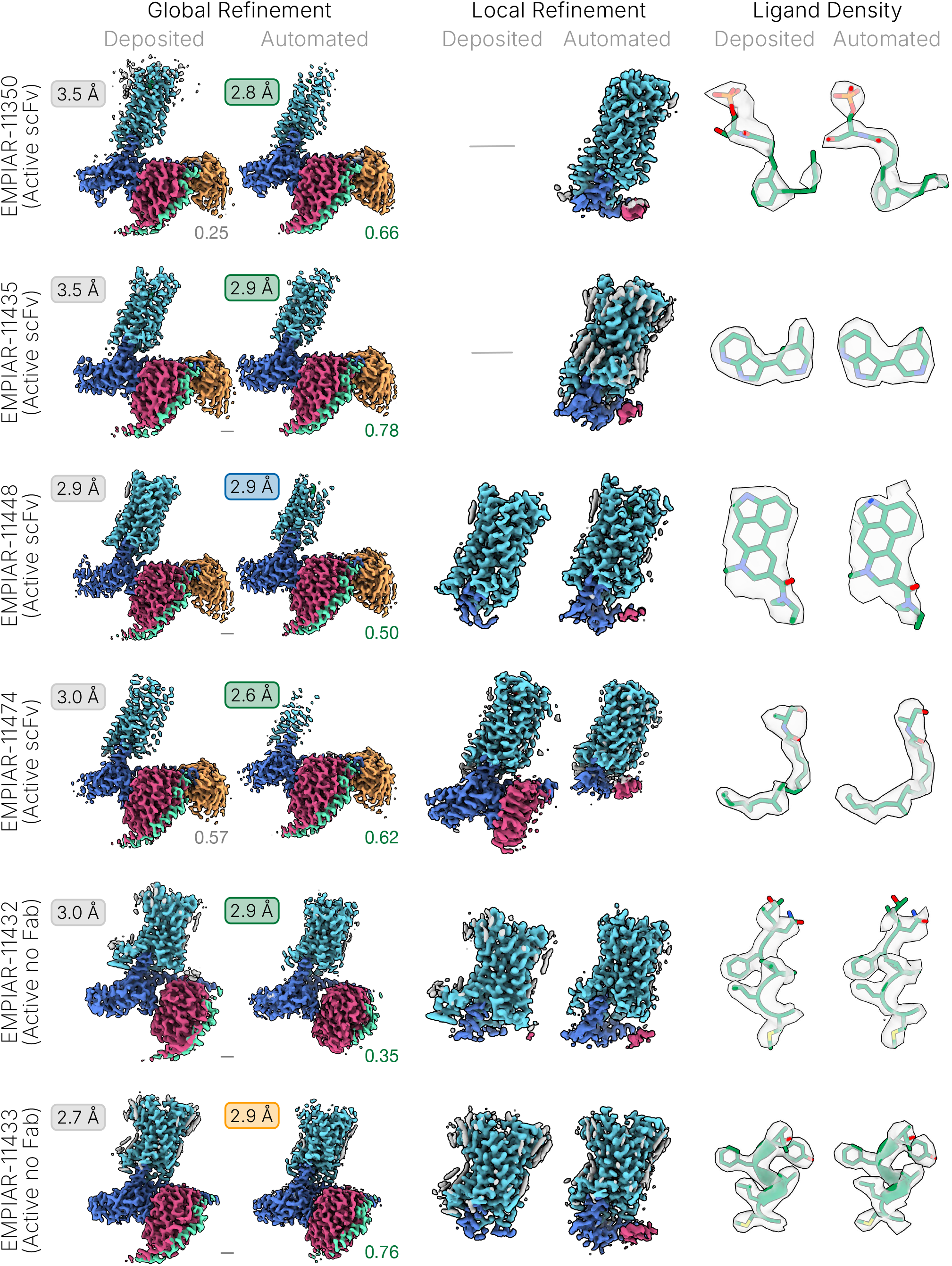
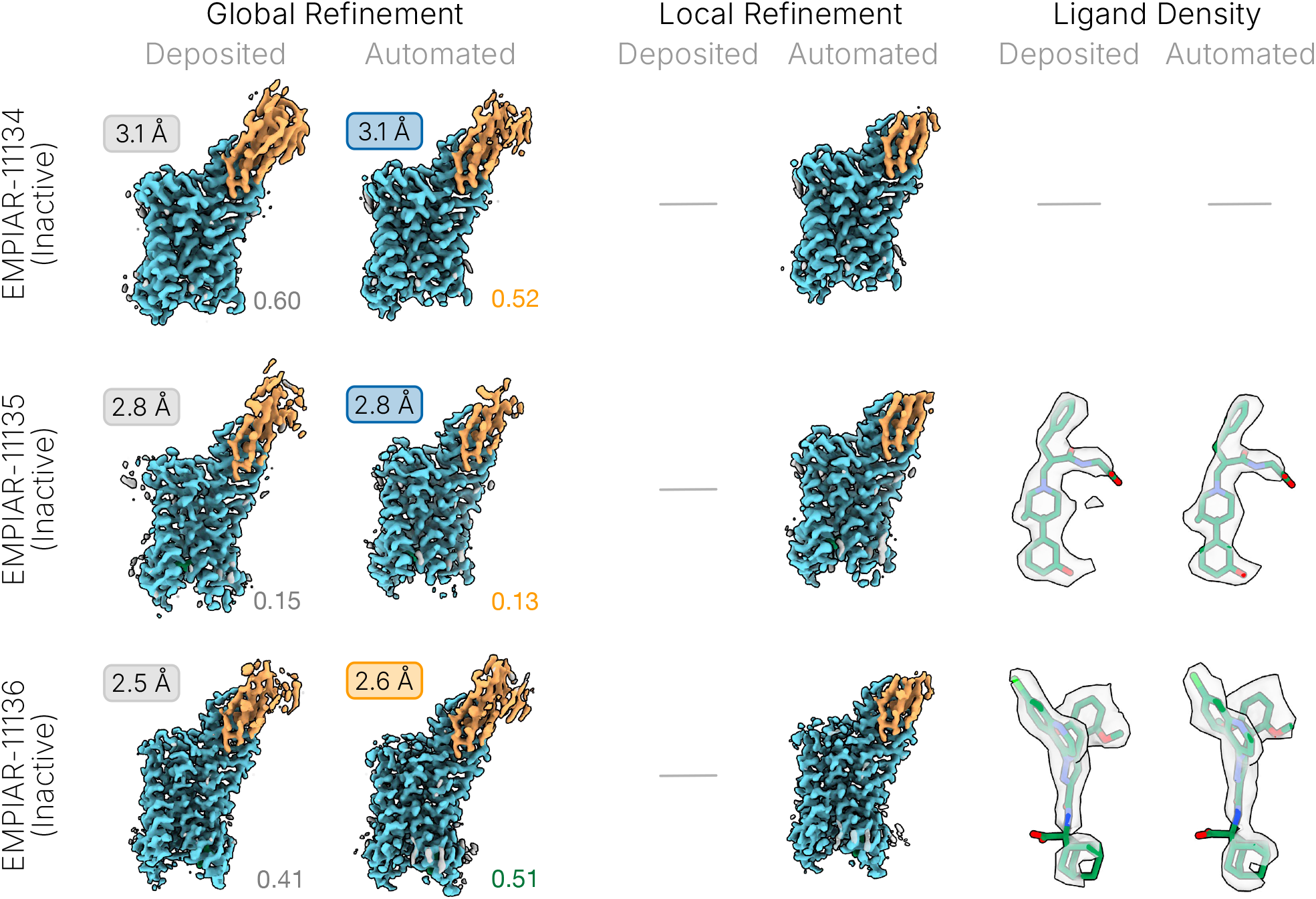
Automated processing yields global, local, and ligand density quality that is equal or better than deposited manual processing across most datasets. Deposited maps and our automated processing maps are shown at similar thresholds for each dataset. For each global map, FSC resolutions (upper left) and cFAR score (lower right) are shown. For automated maps, color scheme denotes improved (green), equivalent (blue), or worse (orange) relative to the manually processed deposited maps. Ligand density is shown with the published atomic model docked for deposited maps, and a re-built atomic model docked for our automated results, to emphasize the improved map quality and interpretability. In some cases, substantially more of the ligand model could be built into the automated density map. For local refinement, dashes represent no deposited local refinement map. For EMPIAR-11119 no ligand density is present in the deposited map. For EMPIAR-11134, no ligand is present in the sample.

### 4.3 Improved Map Quality and Interpretability

For most repeat-target scenarios, the final map quality and interpretability in the region of interest (e.g., ligand binding pocket) are by far the most important metrics of successful data processing. Figure 5 (right column) displays ligand densities from our automated local refinement results in comparison to published deposited maps. In all cases, the map quality of the ligand was similar or improved; and in 10 of 21 cases, density quality was sufficiently improved in terms of improved connectivity and better resolved features to allow for more accurate modeling. In those cases, we remodelled the ligand, and the updated model is shown overlaid with our density in Figure 5. All density maps are rendered using UCSF ChimeraX [41].

Figure 6 showcases three particular examples where our automated results enabled improved model building. For EMPIAR-10288, our map revealed well-defined density for sidechains and loops, allowing modeling of three loops and the N-termini that were not fully built in the original deposited model (PDB-6N4B). For EMPIAR-10855, we were able to resolve a large portion of the ligand along with the previously unmodeled N-terminus. In this case, 2.4M particles remain after automated curation, and it is likely the map can be further improved through extensive 3D classification. A CryoSPARC processing case study exemplified these kinds of improvements on a related dataset, EMPIAR-10853 [42]. Lastly, for EMPIAR-11119, we were able to fully resolve the ligand (PGE2) in the receptor, which was not modeled by the original depositors; they mention that “the extracellular and the ligand binding region of EP3 displayed weak cryo-EM densities” [33]. Furthermore, we were able to model a modest number of backbone and sidechains for the extracellular region.

**Figure 6.**
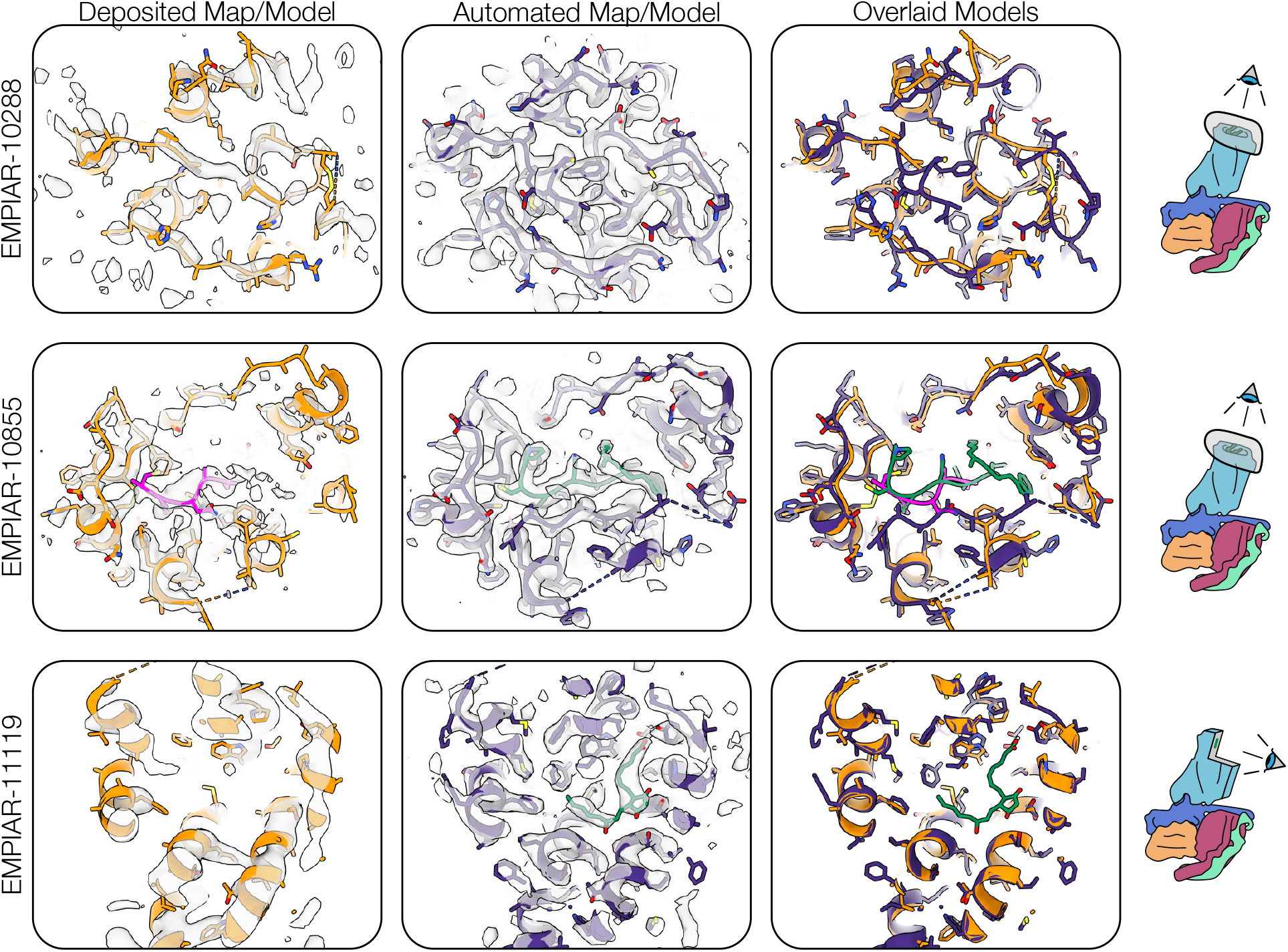
Automated processing yields improved map quality that can enable improved model building. For three examples, the deposited map and model are overlaid and compared with the automated processing map and a re-built atomic model. For EMPIAR-11119, automated results enable fully resolving the ligand (PGE2) in the receptor, which could not be modeled in the original deposited map.

Extracellular loop regions, which are typically flexible and challenging to resolve, are often important for studies of GPCRs because these loops can play a role in binding of orthosteric ligands. Figure 7 displays local refinement maps from our automated processing of all datasets, colored by local resolution to help indicate the relative quality of different parts of each map. The figure shows that after automated processing, the resolution of the receptor domain was relatively high across all datasets, and loop regions were equally well resolved for about half of the cases, while in the other half, the loops were resolved to 3-4 Å. It is likely that the use of local refinement masks more specific to a particular GPCR species (rather than the broad target classes we defined) could result in further improved loop density.

**Figure 7.**
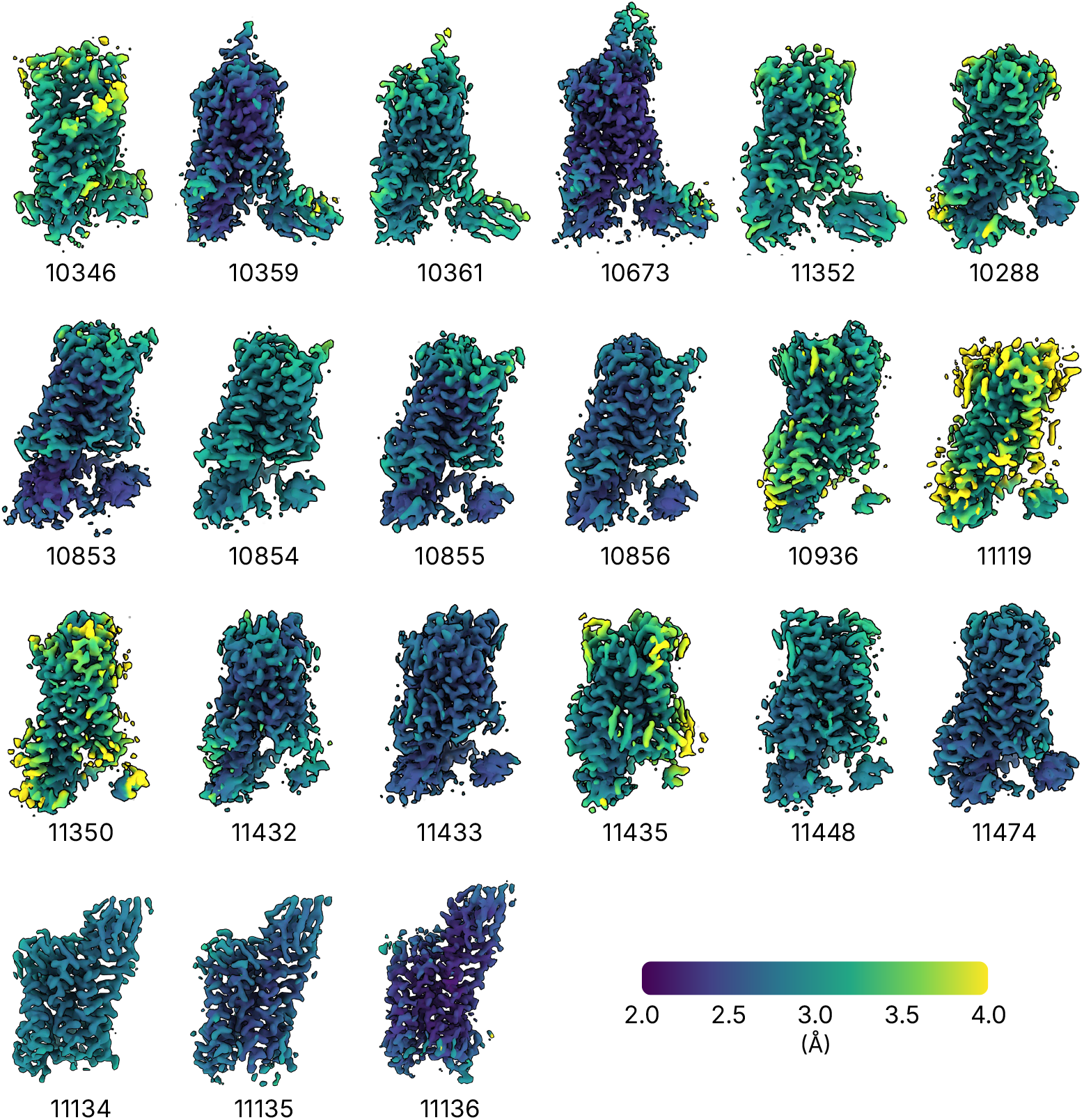
Automated processing yields local refinement maps with intact extracellular loop density in many cases. Extracellular loops are an important feature as they can play a role in ligand binding. Maps are colored by local resolution to indicate the relative quality of different parts of each map.

### 4.4 Robust Particle Curation

Curation of picked particles into a final stack is crucial for achieving high resolution automated results. Our 21 test datasets comprise a wide array of data quality, each with a unique mix of imaging quality, contamination, junk particles, degraded particles, and orientation bias. Being able to achieve successful automation results across these diverse datasets therefore demonstrates that our automation strategy is robust to these variations. Figure 8 displays graphically the relative size of each dataset in terms of number of initial template-picked particle locations, the number of particles retained after micrograph junk detection, automatic inspect picks, decoy classification, and ab-initio reconstruction and heterogeneous refinement to form the final particle set. Notably, the automated workflow is able to successfully curate particle stacks ranging from 1 million up to 12 million initial picks, and the fraction of final retained particles ranges from 2.4% to 28% (with 16% on average). Table S3 further details the number of particles retained after each stage of picking and curation across all datasets.

**Figure 8.**
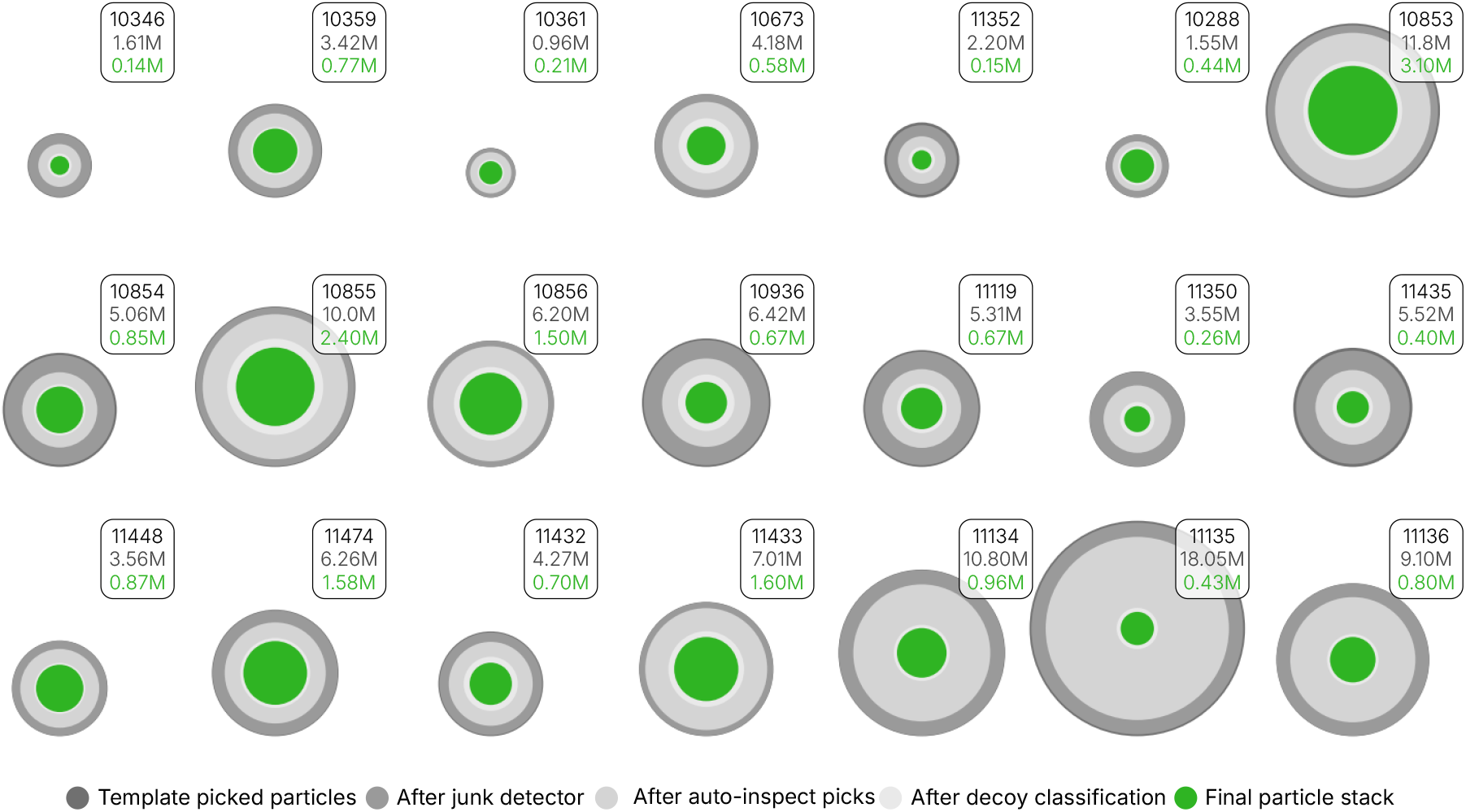
The automated processing workflow is robust to variation in data quality and can isolate high quality particles even when they comprise a small fraction of the dataset. Each dataset is shown as a series of concentric circles with areas proportional to the number of particles remaining at a given stage of processing. The number of template picked particles and number of final particles are shown at the top right for each dataset. The automated workflow is able to successfully curate datasets ranging from 1 to 18 million initial particles, while the fraction of final particles ranges from 2.4 to 28 percent (see Table S3).

### 4.5 Minimal Processing Time and Data Storage Requirements

CryoSPARC includes optimized methods, algorithms and fine-tuned GPU implementations built to minimize time-to-structure. We found that our fully automated end-to-end workflow could be completed within hours on typical hardware. We first processed each test dataset through our complete workflow using only a single workstation with 2 GPUs (NVIDIA RTX 4090) and 32 CPU cores, for which runtimes are reported in Table S4. Processing times varied primarily depending on the number of movies, the pixel size, whether datasets were collected in super-resolution mode, and the number of particles carried through particle curation and refinement. Small datasets (e.g., EMPIAR-10361 containing 1,013 movies) were processed in as little as 9.4 hours while large datasets (e.g., EMPIAR-11135 containing 14,634 movies) took 65.7 hours on a 2 GPU workstation. Across the 21 datasets (on average containing 6400 movies each), the average end-to-end runtime from raw movies to final structure was 34.1 hours. These results demonstrate that even a small 2 GPU workstation can produce rapid turnaround times for real life datasets. However, in practical drug discovery settings where minimizing time-to-structure is key, additional computational resources can be used to further reduce time-to-structure. We repeated the pre-processing and reference based motion correction workflow stages for five of the datasets using a more powerful 8 GPU, 32 CPU server, yielding a 2-3x speedup in those stages and overall speedup of up to 1.7x for the end-to-end workflow on large datasets (Table S4, gray entries). With the correct selection of hardware, it is reasonable to expect that automated processing in CryoSPARC of a typical 6,000 movie GPCR dataset can yield a sub-3Å final map in under 24 hours.

One of the most time-consuming workflow steps is reference based motion correction. Across all datasets, this step takes 30% of the total runtime on average. It is worth noting that this is one of the final processing steps in the workflow and only yields improvements in some cases (Table S2), and therefore can be omitted if necessary to produce further time-to-structure improvements.

It is also worth noting that nearly all of the pre-processing required for our automated workflow can be done during data collection using CryoSPARC Live [6]. A Live Session can be set up with a single click using a configuration profile and will yield on-the-fly pre-processing, micrograph curation, 2D classification, and 3D refinement results that can be used to monitor and diagnose data collection issues. At any point during or at the end of collection, the pre-processed micrographs can be seamlessly connected into a CryoSPARC workspace and the remaining stages of the automated processing workflow can be triggered with one click (see Practical Usage and Automating New Target Classes for details). CryoSPARC Live can be used in this way to reduce time-to-structure by overlapping pre-processing time with data collection, saving an average of 12.5 hours across our test set (Table S4).

Along with processing time, data storage requirements quickly become an important concern for repeat-target processing scenarios. A typical dataset often comprises multiple terabytes of raw data, and the project data generated during processing typically also requires a similar amount of space. For all datasets in this work, we enabled float-16 mode when saving micrographs and particle stacks in order to reduce the data storage footprint. Table S5 contains the breakdown of data storage space used for each dataset, and shows that in general, project data produced by the jobs in these workflows is on average 52% as large as the imported raw data. By streamlining particle curation and eliminating unnecessary exploratory branches, it is possible that automated workflows can reduce unnecessary generation of project data relative to manual processing workflows, in a repeat-target scenario.

## 5 Discussion and Conclusions

The results presented in the previous section demonstrate that our automation strategy is a successful solution to the challenge of automating data processing in the repeat-target scenario. Our automated results are often of superior quality compared to manual processing across a wide range of test datasets. Across the test set of 21 GPCR datasets, the original manual processing was carried out using CryoSPARC [43], RELION [44], cisTEM [45], or a combination (Table S6), requiring multiple stages of manual inspection and intervention and making use of a wide diversity of methods and tools. In contrast, we have shown that our automated workflow can use the same generalized set of steps for all datasets, and can be easily reused for multiple target classes.

Within a target class, only microscope parameters need to be specified for each new dataset in order to commence automated processing, demonstrating that our generalized strategy is robust to substantial variation in data characteristics. Across the test set, the species of the GPCR molecule, type of ligand, microscope, camera, defocus range, ice thicknesses, signal-to-noise ratio, fraction of good particles, and orientation distribution were all variable, but we were still able to achieve high quality results automatically. The results also demonstrate that our workflow generalizes between target classes using only limited prior information, for example between the Active GPCR no-Fab, Nb35, and scFv classes, by changing only the low resolution reference map and masks. In practice, this means that manual processing of a single exemplar dataset provides sufficient information to automate repeat processing within that target class. Furthermore, our results show that the automation strategy can be generalized to inactive GPCRs by changing only reference maps, masks, and the number of decoy classification rounds, despite this class being even smaller and more challenging.

The success of our automation strategy unlocks several benefits for groups attempting to employ single-particle cryo-EM in repeat-target scenarios. First, time-to-structure and throughput can be easily scaled, for example in SBDD or structural genomics settings. Solving multiple structures per day can now become a reality rather than remaining an aspiration. For example, for a homogenous sample where routine grid preparation is established, multiple grids can be prepared with different ligands, and a microscope can be set up to collect several thousand micrographs per grid, taking a few hours each. An automated processing workflow can be launched in CryoSPARC in parallel for each dataset as it is collected, yielding final density maps of quality equal to or better than manual processing, at a throughput matching data collection and a time-to-structure within 24 hours of collection. Secondly, automated workflows in CryoSPARC can improve results while avoiding user bias, mistakes, and variability. As we have shown, for 17 of 21 datasets, automated results yield equal or better resolution and map quality, and for 10 of 21 datasets, automated results yield improved modeling of the ligand, ligand binding pocket, side chain positioning and/or extracellular loops. Consistent, algorithm-driven automated decisions ensure reproducibility across datasets and users, and substantially lower the barrier for new cryo-EM scientists to achieve results that can drive downstream decision-making. Thirdly, by automating routine processing in repeat-target scenarios, researchers can reallocate time and computational resources toward more biologically meaningful questions such as exploring conformational heterogeneity, flexible regions of target molecules, or subtle features such as partial ligand occupancy.

The automation strategy presented in this work, while robust and generalizable, does have limitations. First, automated processing, like manual processing, cannot produce good results from a dataset that is fundamentally limited by poor signal-to-noise or data quality issues. These limitations are most prevalent with smaller particles, such as inactive GPCRs. The three inactive GPCR datasets processed in this work were of high quality and contained a well behaved sample; it should be noted that users working on new datasets may be unable to achieve similar results on small targets depending on sample and data quality. Secondly, targets that are highly flexible may pose an additional challenge for our automated strategy since it relies on using a low resolution reference map. The GPCR test cases in this work contain some flexibility especially in the transmembrane domain, but the static nature of the G-protein complex and presence of the micelle ensure that comparisons against a static reference map are reasonable. Optimization of reference maps may be necessary for extremely flexible targets or targets without a micelle. Finally and most importantly, our automation workflow does not yet attempt to address complex sample heterogeneity, other than curating out broken and junk particles. Extensions of the workflow to cases with discrete heterogeneity using multiple reference maps may be simple, while dealing with continuous heterogeneity remains future work.

All of the methods and tools required to set up an automated workflow for a new target class are already available in CryoSPARC v4.7.1. The next section provides detailed instructions and links to downloadable and importable CryoSPARC Workflow files.

## 6 Practical Usage and Automating New Target Classes

Users can make use of our automation strategy in practice immediately.

At cryosparc.com/automated-workflows we provide instructions and downloadable links for CryoSPARC Workflow JSON files and required inputs which can be imported into CryoSPARC v4.7.1+ and used to process GPCR datasets like the ones we have tested. The Workflow files can be used directly, or as a starting point when automating processing for new target classes. Two versions of the Workflow file are available: one that processes raw movies directly from import, and another that processes micrographs that are generated after pre-processing in CryoSPARC Live.

Performing automated processing for a new target class is straightforward in CryoSPARC. Our strategy can be applied to any type of target, including but not limited to, membrane proteins, soluble proteins, nucleic acid samples, nucleoprotein targets, small proteins and large complexes. As described in Section 3 and Figure 2, it is necessary to gather and set the target-class level inputs (a low-resolution reference map and masks, junk volumes, particle diameter, and particle separation distance) and optionally the workflow level inputs (number of rounds of decoy classification, 2D and 3D selection thresholds). These can then remain fixed for each dataset in the class.

Below is a simple protocol for setting up a new automation Workflow for a new target class, by manually processing an exemplar dataset and then saving the resulting processing steps as a new Workflow. Once this setup is done, the Workflow can be saved and re-used in a single click for new datasets.

1. Decide on a definition of the target class (e.g., GLP-1 receptor with Nb6 nanobody)
2. Choose an exemplar cryo-EM dataset from the target class to use for instantiating the workflow.
3. Find a reference density map for the target class, for example from a previous manual processing of the exemplar dataset, from EMDB, or from structure prediction tools. This reference should be as similar as possible to the target class, but only needs to be approximately 15Å in resolution. Care should be taken to ensure that box sizes are appropriate.
4. As a starting point, import the GPCR automation Workflow JSON file (available at LINK) into CryoSPARC v4.7.1+. Apply the workflow in a new project, but do not queue all the jobs.
  a. Modify the Import Movies job to import the exemplar dataset and associated microscope parameters including exposure groups.
  b. Modify the Import Volumes jobs to import the reference density for this target class.
5. Run each job in the Workflow to process the exemplar dataset. At each of the following points, inspect results and re-run jobs to set parameters, as appropriate, before proceeding to the next job:
  a. Blob picking: adjust particle diameter and particle separation distance for blob picking, and visualize picks using Inspect Particle Picks to confirm.
  b. Reference Based Auto Select 2D: adjust selection thresholds if necessary so that only good classes are selected.
  c. Template picking: update parameters using the values from Blob picking.
  d. Decoy classification: import existing junk volumes from a similar target class, or produce new junk volumes by running Ab-initio Reconstruction and terminating it early or manually editing and importing volumes. Add additional rounds of decoy classification if a single round has not sufficiently curated particles.
  e. Reference Based Auto Select 3D: adjust selection thresholds if necessary.
  f. Local Refinement: produce a local mask around the region of interest using Volume Tools. Masks should have adequate dilation (3-5Å) and a very soft edge (3x dilation distance).
6. In addition to the above, if prior processing experience with the target class is available, any other processing parameters (e.g. in exposure curation, 2D classification, refinements, etc) can be modified to optimize for the target class.
7. Once the Workflow is working for the exemplar dataset, select all the jobs and save as a new Workflow.
8. The new Workflow can now be re-used in a single click for fully automated processing of new datasets from the target class.

## 7 Methods and Extended Results

The following sections describe each stage of processing in the automated workflow presented in Section 3 and Figure 2, including the development of new methods and tools in CryoSPARC that enable automation as well as the experiments we carried out to determine the appropriate strategy for automating each stage of processing. The final workflow is comprised of the following stages and CryoSPARC jobs:

1. Reference, mask, and junk volume imports
  a. Import Volumes (low-resolution reference map, local refinement mask, junk volumes)
2. Pre-processing
  a. Import Movies (using per-dataset microscope parameters)
  b. Patch Motion Correction (float16 and Output F-crop factor = 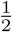 for super-res movies)
  c. Patch CTF Estimation (default parameters)
  d. Curate Exposures (CTF fit resolution *<* 4Å and Total motion *<* 200 px)
  e. Micrograph Denoiser (default parameters)
  f. Exposure Group Utilities (splitting based on filenames using regex)
  g. Micrograph Junk Detector (default parameters, producing junk masks)
3. Particle Picking
  a. Blob Picker on 400 denoised micrographs (diameters and separation distance based on class)
  b. Micrograph Junk Detector (particle rejection only based on junk masks)
  c. Inspect Particle Picks (auto-cluster mode)
  d. Extract from Micrographs (300 px, Fourier-crop to 100 px)
  e. 2D Classification (100 classes, 150Å circular mask)
  f. Reference Based Auto Select 2D (exclude classes worse than 10Å, 150Å circular mask)
  g. Template Picker on all denoised micrographs (diameters and separation distance based on class)
  h. Micrograph Junk Detector (particle rejection only based on junk masks)
  i. Inspect Particle Picks (auto-cluster mode)
  j. Extract from Micrographs (box size matching reference, Fourier-crop to 128 px)
4. Particle Curation
  a. Rounds of Decoy Classification, via:
    i. Heterogeneous Refinement (batch size 3000 per class, using reference and junk volumes as input)
    ii. Subset Particles by Statistic (probability 0.8 or higher in the target class)
  b. Remove Duplicate Particles (separation distance 80Å)
  c. Extract from Micrographs (box size matching reference, Fourier-crop to 300 px)
  d. Ab-initio Reconstruction (5-class, 100,000 particles, max resolution 10Å)
  e. Heterogeneous Refinement (batch size 3000 per class, 12Å initial lowpass filter, all Ab-initio input volumes and all particles)
  f. Reference Based Auto Select 3D (90% or 95% of the best resolution)
5. Final Refinements
  a. Non-Uniform Refinement x4 (each with different CTF refinement parameters)
  b. Select Volume (selecting top refinement by FSC resolution)
  c. Align 3D Maps (aligning selected refinement to reference)
  d. Local Refinement (4 degrees gaussian prior over rotation, 1.4Å prior over shifts, 8Å initial lowpass filter, using mask around region of interest)
  e. Reference Based Motion Correction (using output of Select Volume, float16, Fourier-crop to 300 px)
  f. Non-Uniform Refinement (using output of Reference-based Motion Correction)

This final workflow is captured by the Workflow JSON files that can be downloaded at cryosparc.com/automated-workflows and imported into a CryoSPARC v4.7.1+ instance.

### 7.1 Pre-processing

Each dataset was imported into CryoSPARC via an Import Movies job. For project organization, each import was done in a separate CryoSPARC workspace within a singular project. Microscope parameters for each dataset are detailed in Table S1. Briefly, pixel sizes ranged from 0.675 to 0.91 Å/px, total exposure dose ranged from 44 to 80 *e*^−^/Å^2^, and data from both 200 keV and 300 keV microscopes were used. Dataset metadata was obtained from the respective EMPIAR deposition or corresponding publication.

Patch Motion Correction and Patch CTF Estimation were used to pre-process each of the 21 datasets, with the exception that for datasets that were collected in super-resolution mode, the Output F-crop factor was set to 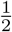 in Patch Motion Correction.

In typical manual processing, the Manually Curate Exposures job can be used to curate micrographs visually by interactively setting thresholds on motion or CTF diagnostic outputs, thereby discarding micrographs that are unusable. To enable automation, these thresholds can be pre-set, but need to be generalizable across datasets within the target class. Through experimentation on eight datasets, we determined that setting two thresholds, namely the CTF Fit Resolution (*<* 4Å) and Total Full Frame Motion (*<* 200 pixels), were sufficient to remove the majority of very poor micrographs that contained no salvageable particles. In the final results running the workflow across all 21 datasets, these thresholds rejected on average 7% of the micrographs from each dataset, with the exception of EMPIAR-11432 where 48% of micrographs were rejected due to poor CTF fits from overfocused images present in the data set. Once micrographs were curated, exposure groups were assigned based on exposure group information present in the filename conventions produced by data collection software.

### 7.2 Particle Picking

Particle picking (and inspection) can be difficult for challenging targets like GPCRs, and existing particle picking methods have tradeoffs that determine their applicability in the automation scenario. Simple sliding-window pickers, such as blob and template-based pickers in CryoSPARC, need only minimal information about the target particle, but can have relatively poor precision that can lead to a large fraction of picked particles needing to be sorted and discarded during particle curation. Trainable neural network pickers, such as Topaz, CrYOLO and others [7, 46, 47, 48], can be trained to improve precision, but require high quality training data that is typically manually curated and potentially dataset-specific. In addition, if rare views or conformations are missing from the training data, neural networks can learn to emphasize this bias. Particle picking methods in general are also susceptible to inclusion of false positives due to the presence of junk features (e.g., carbon edges, ice crystals, cracks, ethane contamination, etc.) and due to inherent noise within micrographs, especially in empty ice regions.

We aimed to find a picking strategy that could limit the inclusion of junk particles regardless of their prevalence in the dataset, while attempting to retain all possible true particles to avoid missing rare views or producing orientation bias. We considered both sliding-window and neural network pickers. Through development of new methods and tools that improve the quality of picks from sliding-window pickers, we were able to produce a picking strategy that is robust to junk, requires no training, and can be easily generalized to new data from the target class. This strategy involves performing blob picking on a subset of micrographs, 2D classification of those picks, selection of 2D classes, and template picking using those 2D classes on the full dataset. The strategy employs four major new developments: junk regions of micrographs are automatically segmented and avoided using a trained neural network model, picking is done on denoised micrographs which substantially improves accuracy, particle picking scores are automatically filtered using a clustering method, and 2D classes are automatically selected through comparison to the low-resolution 3D reference map belonging to the target class. Experimentally, we found that this strategy worked as well or better than a trained neural network picker (Topaz), without the requirement for labelled data or neural network training. The following sections describe the new developments that enable the strategy, and the details of the final workflow.

#### 7.2.1 Micrograph Junk Detector

Although curating exposures by CTF Fit Resolution and Total Motion can remove micrographs that are entirely unusable, in most datasets the remaining micrographs still contain regions of junk which should be avoided in order to produce the best particle picks. Such micrograph-level junk generally appears in a few common categories across datasets: carbon or gold edges, crystalline ice, ethane contamination, cracks, etc. To completely automate the avoidance of junk in micrographs, we trained a neural network junk detector model on an internal collection of manually segmented labelled micrographs. The result is a new Micrograph Junk Detector job type in CryoSPARC (released in version v4.7) that includes a pretrained network that is able to label and segment micrographs fully automatically with three categories of junk: carbon or gold edges, intrinsic ice defects (e.g., cracks, non-amorphous ice, etc.), or extrinsic ice defect or contaminant (e.g. transfer ice, ethane contamination, etc.). The job produces junk masks for each micrograph, and can use these to reject particle picks that are too close to junk regions. It also produces micrographand dataset-level statistics about the presence of junk that can be useful for further micrograph curation and diagnosis of upstream issues in data collection, sample preparation, or sample handling. During development, we tested the Micrograph Junk Detector across a variety of datasets outside of the training set, across microscope, camera, and sample types. The quality of junk masks and statistics it produces are pictured in Figure 9.

**Figure 9.**
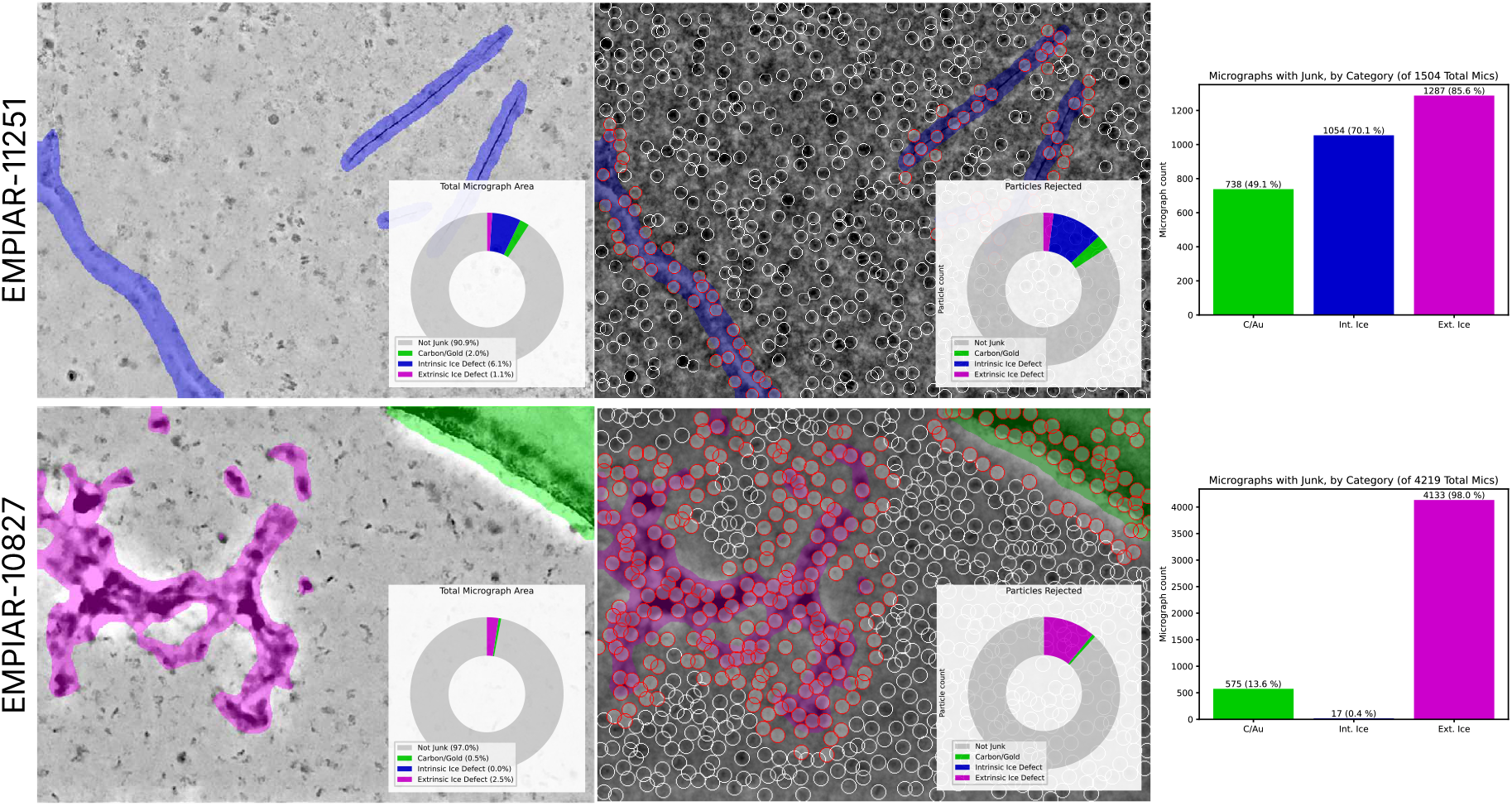
CryoSPARC’s Micrograph Junk Detector (released in v4.7) uses a pre-trained neural network to automatically label and segment junk regions of micrographs and avoid these areas during particle picking. It produces labels in three categories: i) carbon or gold edges, ii) intrinsic ice defects (e.g. cracks, non-amorphous ice, etc.), and iii) extrinsic ice defect or contaminant (e.g. transfer ice, ethane contamination, etc.) The job produces junk masks for each micrograph that are used to reject particle picks close to junk and also to produce micrographand dataset-level statistics.

In our automated workflow, we first ran the Micrograph Junk Detector job during the pre-processing stage, generating junk masks for each micrograph. After picking particles using the blob picker on a subset of micrographs, the Micrograph Junk Detector job was used again; when particle locations and micrographs existing with junk masks are provided as input, the job simply filters the particles by rejecting any that are within a specified distance of junk. The job was used once more after template picking on the entire dataset.

Across our 21 GPCR test datasets, the junk detector removed on average 4% of template-picked initial particles with a maximum of 14% of picks being removed from EMPIAR-11352. These picks were often on ethane or grid support material that are detrimental to early data processing steps due to their high signal content that can throw off 2D and 3D reconstructions.

#### 7.2.2 Micrograph Denoiser

Single particle cryo-EM micrographs are collected with limited electron exposure dose to limit sample radiation damage, yielding a high level of irreducible (i.e., truly random) electron shot noise in collected images. In addition, the micrographs also contain a large degree of structured noise as a result of contaminants, solvent molecules, denatured protein, and uncorrected optical aberrations. For smaller and more challenging particles, these noise sources can dominate micrograph images and are part of the reason that particle picking can be highly challenging. During data collection, exposures are recorded as movies, with multiple frames captured per field of view. These multiple frames can be partitioned, for example into even and odd frames, to produce two images that contain the same signal and structured noise, but different measurements of shot noise. Pairs of such images can be used for the *noise2noise* [49] training setup whereby a neural network is trained to denoise one image to match its pair, and vice versa. This approach has been used by several groups to create denoisers for single particle cryo-EM data [7, 50, 51, 52]. Ideally, repeated particle signal stands out during the training process and is emphasized, while noise sources are attenuated.

We developed a new *noise2noise*-based method for micrograph denoising that substantially improves the visual quality of denoised micrographs compared to previous methods by making use of an improved neural network architecture and additional training-time priors on smoothness of the output image. The new method, available in CryoSPARC as the Micrograph Denoiser job (released in v4.5) enables very fast training (less than 5 minutes for a new dataset) and inference (over 300 micrographs per minute), requires only 100 training micrographs per dataset, and is fully automatic with no labels or adjustment of parameters needed. The job can be run during pre-processing, immediately after CTF estimation. The resulting denoised micrographs (examples shown in Figure 10) have far stronger contrast for particles, and in many cases individual particles become visible. This improved contrast compared to the background improves the accuracy of particle picking with sliding-window (i.e., blob and template) pickers, which can now more clearly distinguish true particle candidates from empty regions. Furthermore, the denoiser causes micrographs of differing contrast and defocus levels to become more visually similar, reducing the variation in particle picking scores that can otherwise result. This change makes it possible to make consistent selections of high quality particles based on picking scores across micrographs and across datasets and is crucial for our automated workflow.

**Figure 10.**
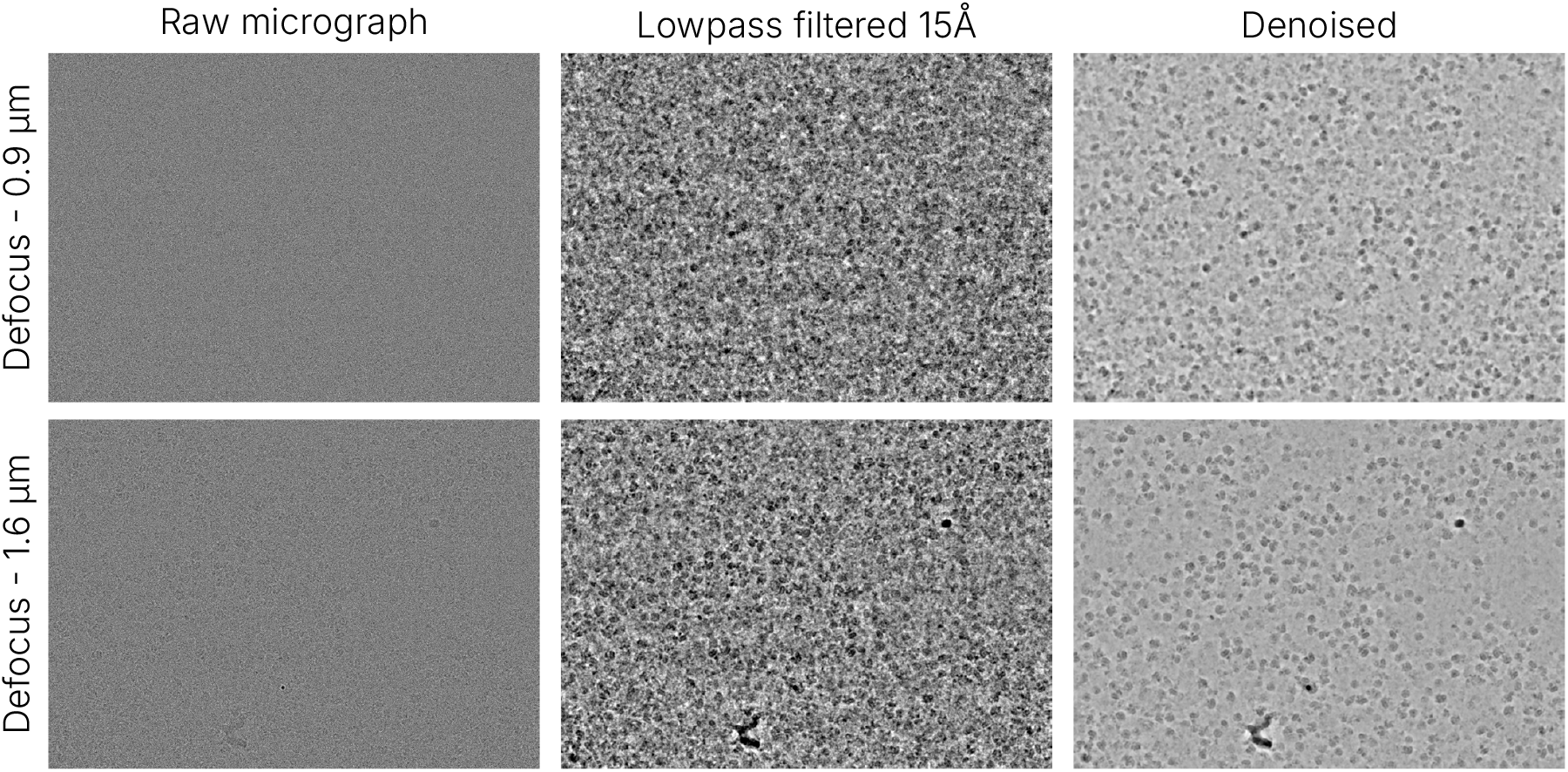
CryoSPARC’s Micrograph Denoiser (released in v4.5) is a *noise2noise*-based method that removes noise and produces improved contrast of individual particles. It trains quickly, taking less than 5 minutes for a new dataset with only 100 training micrographs required, and can be run on new micrographs at over 300 micrographs/minute. Denoised micrographs have improved signal for particles allowing sliding-window (i.e., blob and template pickers) to more clearly distinguish true particle candidates from empty ice regions. Denoising also causes micrographs from different defocus levels to appear similar and to produce comparable particle picking scores.

#### 7.2.3 Inspect Particle Picks Auto Cluster Mode

In a typical manual workflow, a user would sort particles to remove false positive picks (empty ice, high contrast, overlapping or aggregated particles) using the Inspect Particle Picks interactive job. In this job, a user reviews multiple micrographs across a range of defocus values and manually sets thresholds for Power score and Normalized Cross Correlation (NCC) score by examining histogram values and particle locations. As mentioned in the previous sections, the new Micrograph Junk Detector automatically removes particles in junk regions, and the Micrograph Denoiser improves picking accuracy and yields consistent picking scores across micrographs and datasets. During development, we noticed that with these upstream improvements in place, the histogram of particle picking scores across example datasets became more structured, often with clear clusters corresponding to false positive picks in empty ice regions, reasonable candidate particles, and high-contrast picks typically featuring multiple occluding or aggregated particles. We therefore developed a new auto cluster mode for the Inspect Particle Picks job (released in CryoSPARC v4.6) that separates particles into these clusters, and automatically selects the cluster of reasonable candidate particles for downstream processing (Figure 11). This job is used to filter both blob picked particles and template picked particles, and enables fully automated picking without any manual intervention.

**Figure 11.**
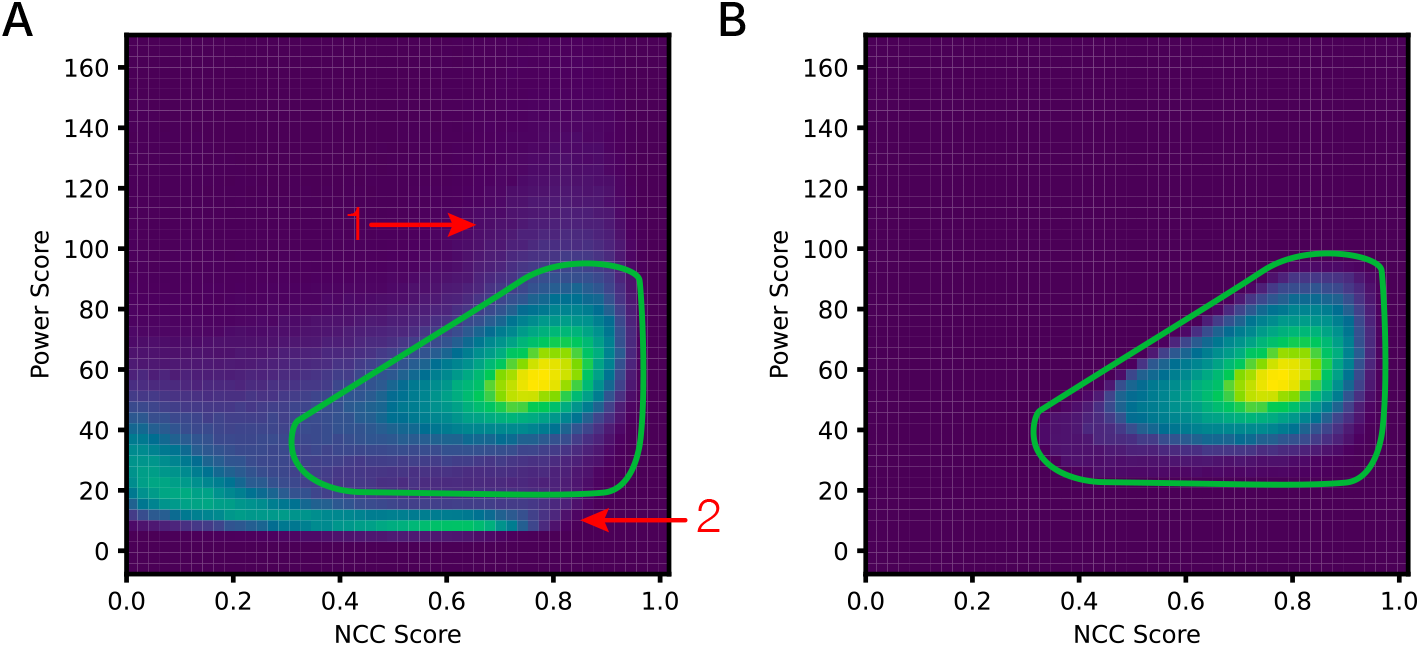
Auto cluster mode in Inspect Particle Picks job (released in v4.6) clusters particle picking scores and automatically selects the cluster of reasonable candidate particle picks (outlined, green), separating those from occluded or aggregated particles (1) and empty ice regions (2).

We compared Inspect Particle Picks auto cluster mode to manual threshold selection for template picks from five datasets, and found that it resulted in very similar final results (Table S7). Across the 21 GPCR test datasets, we found that when picking on denoised micrographs, most reasonable particle picks ended up having a power score of 50-75 while picks over empty ice had power scores of 0-20, and auto cluster mode removed between 15 and 50% of particles.

#### 7.2.4 Reference Based Auto Select 2D

In developing our automated workflow, we initially experimented with blob picking alone and found that, with the addition of the three preceding improvements, the quality of automatically produced blob picks on challenging GPCR datasets was substantially improved relative to a baseline of applying blob picking to raw micrographs and only filtering the particles manually. However, we found that many particle picks were slightly off-center, and some particles were missed while others were picked twice. These are known limitations of blob picking, which does not take any information about the shape of the particle into account. We therefore considered the approach of classifying blob picked particles in 2D, and then using the resulting 2D classes for template picking on denoised micrographs. We ran 2D classification with 100 classes. In the resulting 2D classes, typically between 5 and 20 classes contained the target particle. While such classes can easily be selected by eye, an automated workflow requires the selection to be done automatically.

Several groups have worked on automatic 2D class selection methods using neural networks trained on examples of good and bad 2D classes labelled by expert users [16, 8, 13, 17]. These methods typically use a fixed pre-trained neural network, and the quality of their selection depends on the presence of the target class in the training data that was used. We opted to develop a simple but more robust method for selection of 2D classes by making use of the low-resolution 3D reference map that defines the target class. The new method aligns and compares each 2D class to the 3D reference, and produces two matching scores that can be used to separate classes containing the target from junk and contaminants. Importantly, we developed this method with the goal of selecting 2D classes that would suffice for re-picking with the template picker, rather than for curating particles in 2D, as we found that 3D curation produces improved results and does not lead to the degree of orientation bias that can easily be caused by accidentally excluding views during 2D curation.

The Reference Based Auto Select 2D (RBAS 2D) job (released in CryoSPARC v4.5) produces a score using the Sobel image operator that measures the relative level of similarity in shape, and a Correlation score that measures overall density similarity between each 2D class and the 3D reference (Figure 12). These scores are consistent across datasets and references, and we found that across all 21 GPCR test datasets, classes with correlation score greater than 0.9, Sobel score less than 1.5, and estimated 2D class resolution better than 10Å corresponded to good classes. Using these classes for template picking on denoised micrographs yielded well-centered picks with few missing particles or duplicates.

**Figure 12.**
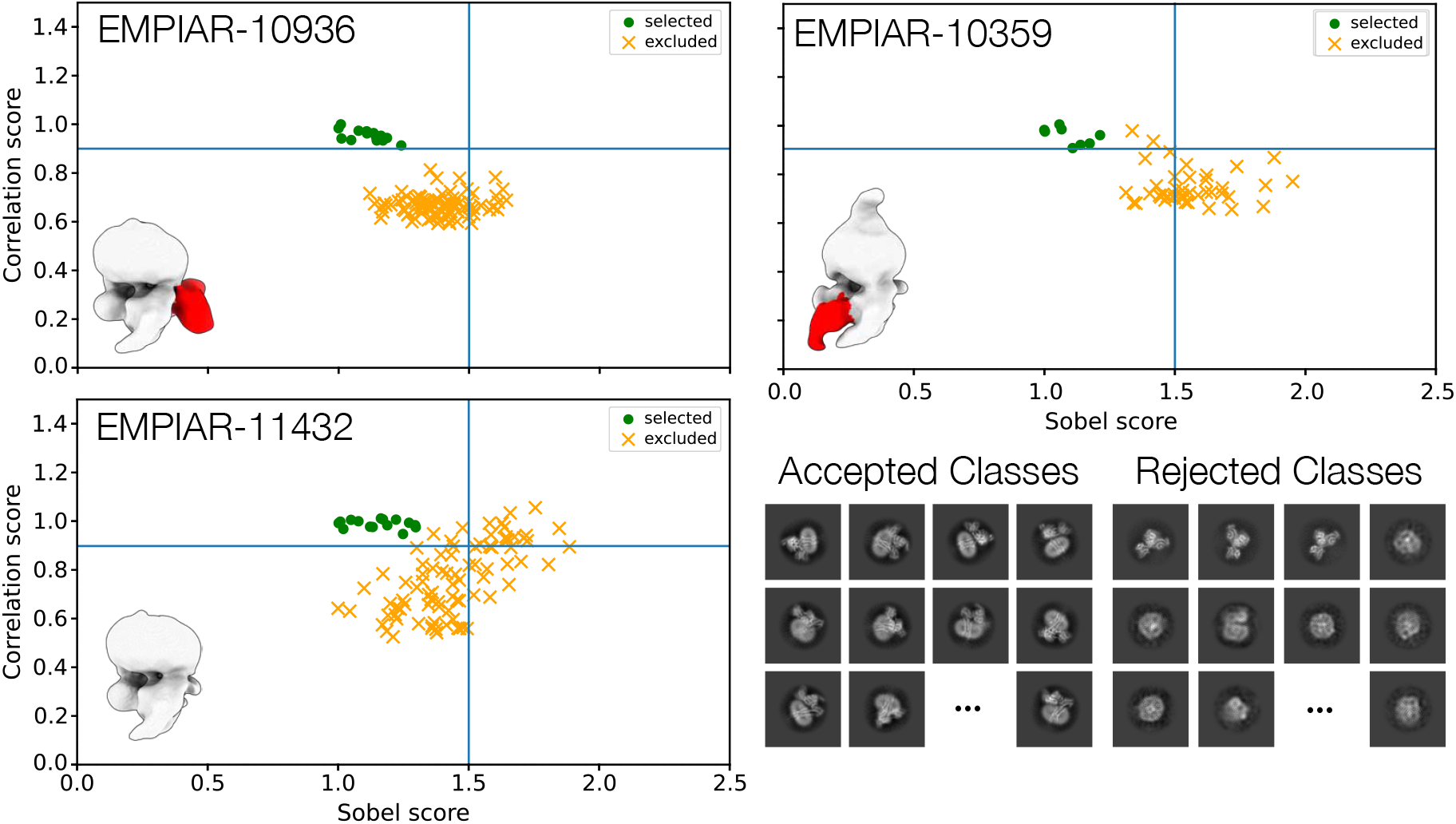
Reference Based Auto Select 2D (RBAS 2D, released in v4.5) automatically selects 2D classes using a 3D reference map. Each 2D class is aligned to the 3D reference map, and two scores are computed: a Sobel match score that measures shape similarity, and a Correlation score that measures overall density agreement. These scores, along with 2D class resolution, separate good 2D classes that contain the target particle from those that do not. Scatter plots of the scores for three datasets are shown, with the 3D reference map inset, default thresholds as blue lines, accepted classes as green circles, and rejected classes as yellow crosses. Examples of accepted and rejected classes are shown in the bottom right panel.

#### 7.2.5 Final Particle Picking Strategy and Comparisons

With the above improvements in place, we were able to construct our final particle picking strategy (Figure 13). An initial set of particle picks were generated by blob picking on 400 denoised micrographs. Blob picking parameters were set as follows: for all active classes, blob picking diameter 110Å (min) to 140Å (max) with separation distance 0.7, and for the inactive class, picking diameter 100Å (min) to 120Å (max) with separation distance 0.7. Picks near junk were rejected using the Micrograph Junk Detector with default parameters. Automatic Inspect Particle Picks was used to remove any remaining false positive picks. The resulting good particles were then extracted at a box size of 300 pixels and Fourier cropped to 100 pixels to save on disk space and processing time. 2D Classification was run with 100 classes, a 150Å circular mask, 40 online-EM iterations, and a batch size of 250 particles per class. Based on the dataset’s target class, the appropriate reference was used for template selection by Reference-based Auto Select 2D, with Sobel score threshold 1.5, Correlation score threshold 0.9, and resolution cutoff 10Å. The selected 2D classes were used for template picking on denoised micrographs for the entire dataset, using a particle diameter of 120Å with separation distance 0.6 for active classes and diameter 110Å with separation distance 0.6 for inactive. The template picked particles were again filtered by the Micrograph Junk Detector and Automatic Inspect Particle Picks. These particles were then carried downstream to particle curation in 3D, with no further 2D classification.

**Figure 13.**
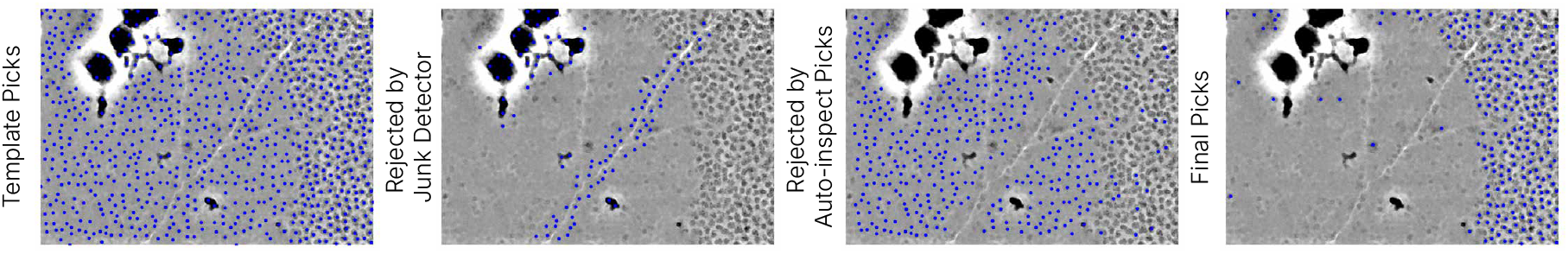
The final particle picking strategy used in our automated workflow produces high-quality particle picks across the various datasets. Blob picks (not shown) on 400 denoised micrographs yield 2D classes that are automatically selected using RBAS 2D (Figure 12). The selected classes are used for template picking on denoised micrographs (left). Picks close to junk are automatically rejected **(center-left)**. Auto cluster mode in Inspect Picks automatically removes empty ice and aggregation picks **(center-right)** yielding a high-quality set of final picks for downstream processing **(right)**.

As a comparison with neural network particle pickers, we compared our picking strategy with Topaz [46] on five GPCR datasets. We used Topaz in two ways. First, for each test dataset, we trained a Topaz model on a subset of the same dataset, and then ran the model on the full dataset. Second, we trained a single Topaz model on four additional datasets outside the test set, and used this joint model on the four test datasets. Positive labelled particles for training data from each dataset were generated by blob picking, removal of junk and empty ice picks, 2D classification, and 2D selection. Across five tested datasets (Table S8), we observed that after particle curation, the quality, resolution, and cFAR scores of maps derived from our strategy were better or equal to those derived from Topaz-picked particles. Furthermore, the individual Topaz models required performing nearly all the same steps as in the automated picking workflow in order to generate training data. In addition, the joint Topaz model would not generalize to new target classes without being trained on each new class.

### 7.3 Particle Curation

Particle curation is perhaps the most critical step in the single particle data processing workflow. Datasets can contain millions of particle candidates at the particle picking stage, but only a small fraction of those, typically less than 10%, are actually used to produce the highest quality final refinement in state-of-theart manual processing workflows. It is not, in general, possible to accurately assess the quality of an individual single particle image due to the very poor signal to noise ratio in cryo-EM data; it is also not possible to assess the impact of a single particle on the final refinement or other downstream analysis, or to perfectly separate true particles from broken or junk particles. In the final curated particle set, however, aggregate particle quality as well as the ratio of intact high-quality particles to residual junk particles have a significant impact on 3D map quality and especially on downstream analysis including separation of conformational or compositional heterogeneity and local refinement of flexible regions.

The particle picking strategy described in the previous section eliminates most micrograph level junk and false positive picks. Of the remaining particles, high-quality particles are those that are intact, in thin ice, and where structured noise sources such as crowding, contaminants, and any layers of denatured protein are minimal so that accurate contrast transfer function (CTF) estimation and pose alignment is possible. The low-quality particles typically are denatured, aggregated, broken, partially formed, or have too much nearby structured noise.

Particle curation, starting from the full set of extracted particles, can be performed using either 2D classification or 3D approaches. 2D classification is in general simpler, but treats particle images as examples from multiple independent 2D density maps, without exploiting the inherent relationships between different viewing directions of the same 3D molecule. Furthermore, 2D classification is run typically with 50-400 classes, which is much smaller than the number of potentially distinct viewing directions of a 3D object. Therefore, it is often the case that rare orientations are inadvertently discarded because they were lumped in with junk classes, unnecessarily inducing orientation bias. In contrast, 3D particle curation ensures all viewing directions are modelled while allowing shared information from common orientations to aid the classification of rare views. Typically during 3D curation, “junk” density maps will form that absorb outlier particles across multiple orientations. For these reasons, we opted to use only 3D curation methods in our automated workflow. 3D curation, which makes use of 3D classification and refinement methods, is substantially more computationally complex than 2D classification given the expanded search over 3D poses and classes that must be carried out for each particle. CryoSPARC is particularly efficient for 3D curation due to its use of specialized algorithms for pose alignment and finely tuned GPU implementations [43].

A typical 3D curation protocol that is widely employed by CryoSPARC users begins with multi-class Ab-Initio Reconstruction to generate a dataset-specific set of initial volumes, each containing either the intact target structure or contaminant junk density. These initial low-resolution volumes are then jointly refined using Heterogeneous Refinement, which allows each volume to progress to higher resolution and improves particle classification accuracy. Particles assigned to high-quality classes are retained for downstream processing and the rest are discarded, and the protocol can optionally be repeated if necessary. This approach can only succeed if Ab-Initio Reconstruction generates at least one intact initial volume containing the target molecule; for datasets with a small fraction of high-quality particles, the likelihood of success decreases, and is particularly unlikely for smaller targets such as GPCRs, where signal-to-noise ratios are inherently lower. In our experiments, we found that using multi-class Ab-Initio Reconstruction immediately after particle picking failed to produce any intact volumes for several datasets, especially inactive GPCRs. An automated workflow based on this approach would fail at this point, so we explored alternative curation strategies.

#### 7.3.1 Decoy Classification

To robustly handle datasets where high-quality particles comprise a small fraction of total particles, we used a 3D curation strategy that is already employed by some CryoSPARC users in the manual processing setting, referred to as decoy classification. Decoy classification starts with a known set of volumes as inputs to a Heterogeneous Refinement job. One of these volumes is an intact low-resolution reference volume of the target molecule, and the rest are junk volumes. During classification, the reference volume attracts intact high-quality particles, while the diverse junk classes help to absorb poor-quality particles. For GPCRs, we produced four junk volumes (see Supplementary Materials for details) and used the 15Å target-class reference volume as inputs. Heterogeneous Refinement further filters the input volumes to 20Å. This strong level of filtering means there is no risk that the reference produces bias in the higher resolution features of the results.

Across the active GPCR datasets, a single round of decoy classification proved to be a straightforward and universally applicable approach for the initial pass of automated 3D curation. For inactive GPCR datasets, we observed that curation results could be improved by using two rounds of decoy classification. To further increase the specificity of classification, after each round we selected only particles that were classified into the intact class with probability exceeding 0.8, using the Subset Particles by Statistic job (released in CryoSPARC v4.7). The optimal number of rounds of decoy classification for a given target class can vary depending on factors such as particle size, expected signal-to-noise ratio (SNR), and the prevalence of junk particles. In most cases, decoy classification eliminated 23-68% of picked particles (Table S3). In our experiments we found that the resulting set of particles had a substantially increased fraction of high-quality particles, but refinements of these particle stacks did not yet produce results similar in quality to manual processing. One reason we suspected for this result was that decoy classification makes use of 3D references and junk volumes that are generic rather than dataset specific.

#### 7.3.2 Ab-initio Reconstruction and Heterogeneous Refinement

We therefore next employed a round of 3D curation using Ab-initio Reconstruction to generate new initial volumes from scratch, followed by Heterogeneous Refinement starting from those volumes. At this stage, starting from a cleaner particle stack produced by decoy classification, we found that across our test datasets, a 5-class Ab-Initio Reconstruction job using a subset of 100,000 particles produced 1-3 intact volumes containing the target molecule, with the remainder of the volumes representing junk particles. The subsequent Heterogeneous Refinement used all particles and all five ab-initio generated volumes, classifying particles into either intact or junk classes. This approach was effective as a final curation step, likely due to the improved dataset-specific references provided by Ab-initio Reconstruction. An added benefit of this stage is that 3D references are not provided as input, further increasing confidence that resulting volumes are not biased by input references.

Ab-initio Reconstruction is not guaranteed to produce a specific number of intact vs. junk volumes, or to produce those volumes in a particular order. For complete automation, it is necessary that the intact volumes be selected so that the corresponding curated particles can automatically be used downstream. To automate this selection, we developed a new tool that can identify suitable classes using either a 3D reference structure, or simple resolution metrics.

#### 7.3.3 Reference Based Auto Select 3D

CryoSPARC v4.5 introduced the Reference Based Auto Select 3D (RBAS 3D) job which provides two modes for selecting volumes and their corresponding particle stacks from a set of inputs. In reference-based mode (the default), input volumes are aligned to the supplied reference volume, and FSC and Correlation scores are computed between them. Volumes with scores higher than specified thresholds are selected for downstream processing. In the simpler resolution-based mode, the best input volume is selected based on resolution, and any other volumes with a resolution that is within a specified percentage are selected as well.

We found that the simple resolution-based mode worked well across both the active and inactive GPCR datasets. For other target classes, reference-based mode may aid in accurately selecting volumes. Figure 14 displays examples of reference-based selection of 3D classes.

**Figure 14.**
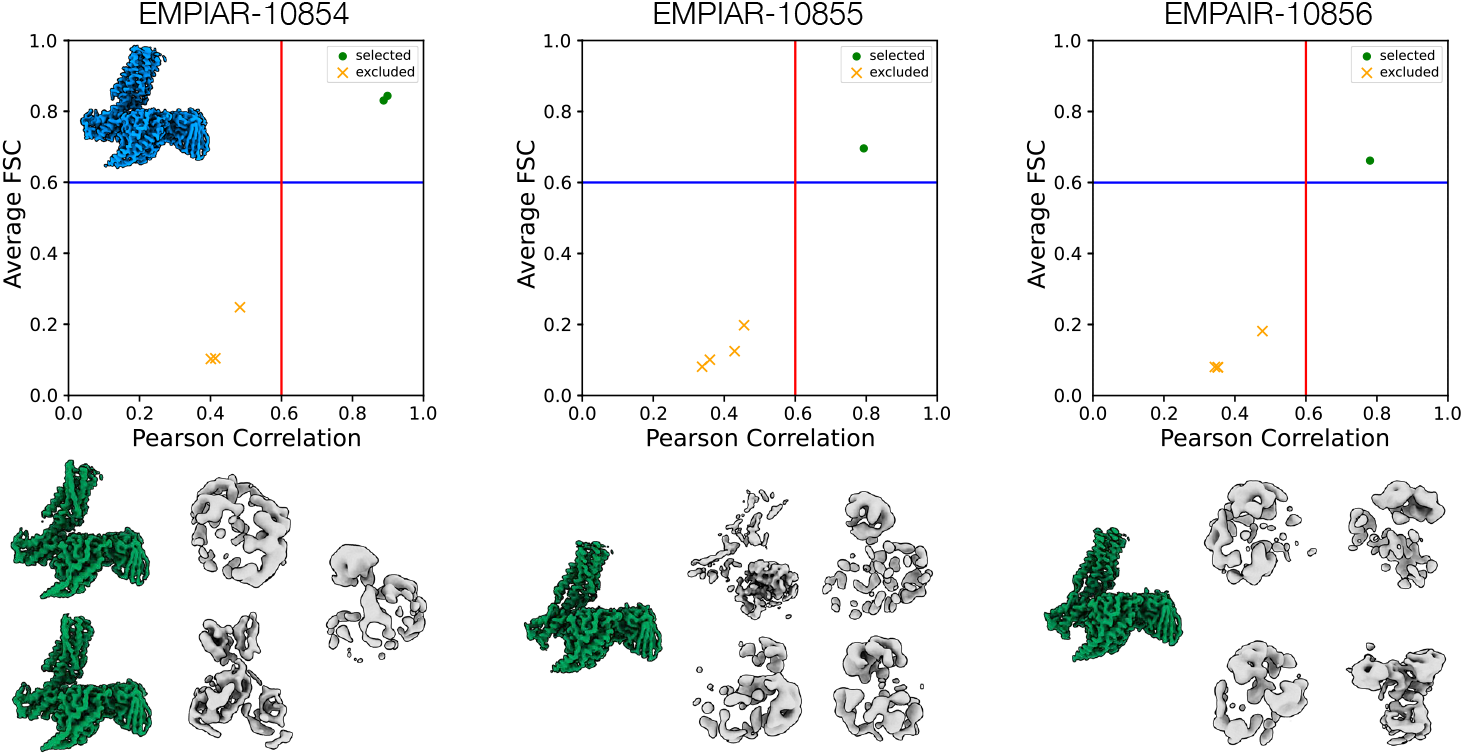
Reference Based Auto Select 3D (RBAS 3D, released in CryoSPARC v4.5) can operate in two modes: in the simpler resolution mode (used in our automated workflow, but not shown here), the best class is selected based on resolution and other classes within a specified percentage of that resolution are selected as well. In reference mode (pictured here, but not used for our automated results), a 3D reference (inset, blue, from EMPIAR-10853) is used. Each 3D class is aligned to the reference, and the FSC and Pearson correlation between the 3D class and the reference are used to select classes. Reference mode can be useful for more challenging targets where separating high quality 3D classes by resolution is difficult. For each of three datasets shown, the scatter plot shows selected 3D classes as green circles and rejected as yellow crosses. The bottom row shows the selected densities in green and rejected in grey.

Our final strategy for particle curation, involving rounds of decoy classification, Ab-initio Reconstruction and Heterogeneous Refinement, and Reference Based Auto Select 3D, was able to robustly curate particles across all 21 datasets in our test set, providing final particle stacks that could be refined to resolutions and map quality matching or exceeding published manual processing results.

### 7.4 Final Refinements

CryoSPARC’s Non-Uniform Refinement [53] is widely used for small membrane proteins such as GPCRs due to the improved results it can often generate by accounting for the spatially variable properties of the particle. For example, it automatically differentiates between intact and rigid protein density, and regions of local flexibility and disorder such as a nanodisc or micelle. In our tests, we found that when starting from curated particles, Non-Uniform Refinement produced high quality and high resolution density maps. We also found that for some datasets, certain combinations of CTF refinement parameters could improve the results, as could optimizing motion trajectories using Reference Based Motion Correction. However, in some cases, these additional steps made results worse. For complete automation, we therefore opted to simply run multiple refinements and then select the refinement that produces the best result.

Furthermore, we found that after global refinement using Non-Uniform refinement, map quality was generally best in the G-protein complex (for active GPCR datasets) but poorer in the transmembrane domain where the ligand binds with clear signs of unresolved continuous flexibility. We therefore added Local Refinement to our automated workflow, by first aligning the global refinement result to the original target-class reference map, and then using a target-class-specific mask over the region of interest. This approach yielded substantially improved density quality in many cases and enabled re-building of the atomic model into the receptor and ligand. The following sections provide details about the steps in the final refinement workflow stage.

#### 7.4.1 CTF Refinements

High resolution cryo-EM structures often require correcting for electron-optical aberrations, aberrations introduced by beam-image shift data collection that are not fully compensated for by data collection software, and microscope misalignments that result in nuanced high-order terms in the Contrast Transfer Function (CTF). These higher order terms (beam tilt, trefoil, spherical aberration, tetrafoil, anisotropic magnification) are estimated from single particle data itself, by refining the corresponding CTF parameters against a high-resolution reference map [54]. In CryoSPARC, either the Global CTF Refinement job can be used to correct for these higher-order aberrations, or the CTF refinement process can be run within iterations of refinement jobs by turning on a parameter. The set of aberrations to refine and correct are normally also user-specified parameters. A particular dataset may benefit from correcting only a subset of the aberrations, while attempting to correct for all may make results worse; typically this occurs when the particles are small and images have poor signal quality. While an expert user could inspect diagnostics from a CTF Refinement job and determine which aberrations to correct or ignore, this decision making is not straightforward in an automated setting. Instead, our workflow runs four Non-Uniform Refinement jobs each with different CTF refinement settings: 3rd order; 3rd and 4th order; 3rd order plus anisotropic magnification; 3rd and 4th order plus anisotropic magnification. In a computing environment with multiple GPUs, these refinements can be parallelized to reduce the total runtime of the workflow. The best of the resulting four refined volumes can be selected automatically using the Select Volume tool described in the next section.

We found that Global CTF Refinement produced an improvement of up to 0.2Å across our test set. In addition to Global CTF Refinement, we also tested whether refining per-particle defocus values using Local CTF Refinement would be beneficial. Across the GPCR test set, we did not find any improvement from adding this step, likely due to the relatively thin ice that these datasets exhibit.

#### 7.4.2 Select Volume

To simplify selection between multiple refinement results, we created the Select Volume job (released in CryoSPARC v4.7) that takes in a number of refinement results including both volumes and particle stacks, and outputs the result that has the highest overall gold standard FSC curve, measured based on the area under the FSC curve. This job simplifies automated selection between branches of processing in any scenario.

#### 7.4.3 Reference Based Motion Correction

Alongside refinement of CTF parameters, refinement of particle motion trajectories and dose weights using a reference volume [55] can produce improved map resolution and quality for some datasets, while making results worse for others. In CryoSPARC, the Reference Based Motion Correction job can be run after a refinement, and the resulting particles run through refinement once more to determine if there was an improvement. In our workflow, we performed this step after the Select Volume job so that particles with optimized CTF refinement parameters were used. A subsequent Non-Uniform Refinement (without further CTF refinement) produced a new refined map reflecting the motion and dose weighting optimizations, and Select Volume was again used to determine whether these changes produced an improvement.

Across our test set, Reference Based Motion Correction made up to 0.2Å improvement in resolution on EMPIAR-11352. Six other datasets improved by 0.1Å. Given the additional processing time and storage costs of running this step, it may be desirable to omit this step from workflows for other target classes.

#### 7.4.4 Orientation Diagnostics

CryoSPARC refinement jobs produce diagnostic plots that assess the level of anisotropy in the refined 3D density map that is a result of the orientation distribution of the particles. The level of anisotropy is summarized in the cFAR (conical Fourier Area Ratio) score, which measures the ratio between the best and worst direction in terms of conical FSC curves [40, 56]. The Orientation Diagnostics job in CryoSPARC can also be used as a standalone tool to produce a larger set of diagnostics.

#### 7.4.5 Align 3D Maps and Local Refinement

Local refinement of particle poses, focused using a mask on the region of interest with the 3D map, can substantially improve the quality of density especially for particles with continuous heterogeneity and flexibility between parts. The choice of mask is important, as a mask that is too small can yield poor quality artefactual reconstructions, while a mask that is too large reduces the ability to capture local flexibility. Once an appropriate mask is created, it can be reused across datasets within a target class, as long as the mask is in register with the 3D density map to which it will be applied. In our automation workflow, the local resolution mask is provided as a target-class level input and is in register with the input low-resolution reference map. However, our final particle curation step uses Ab-initio Reconstruction to produce initial volumes from scratch for the specific dataset being processed, and Ab-initio maps can be arbitrarily oriented within the box due to random initialization. Subsequent refinements will therefore also be arbitrarily aligned. To bring refinement results into register with the low-resolution map and local refinement mask, we use the Align 3D Maps job (released in CryoSPARC v4.5). This job compares each input map to a reference map, and computes the optimal 3D rotation and shift to cause them to match. The job applies the rotation to the input maps and their corresponding particle orientations.

For our GPCR test datasets, we produced a local refinement mask for each target class (see Supplementary Materials) to cover the transmembrane domain. Local Refinement was run with 4 degrees standard deviation of rotation, 1.4Å standard deviation of shift, 12 degree rotation search extent, and 4.2Å shift search extent. Across the datasets, locally refined maps displayed improved density quality in the transmembrane domain and especially in the ligand binding region.

## Data and Software Availability

All data processing in this work was carried out using CryoSPARC v4.7.1, available at `cryosparc.com`. At cryosparc.com/automated-workflows we provide instructions and downloadable links for CryoSPARC Workflow JSON files and required inputs which can be imported into CryoSPARC v4.7.1+ and used to replicate our results or extend automated processing to new target classes. All final density maps produced in this work are available upon request. All raw data was downloaded from EMPIAR [23]; see accession codes in Section 2.

### Acknowledgments

We thank the entire team at Structura Biotechnology Inc. that designs, develops, distributes, maintains and supports the CryoSPARC software system with which this project was implemented and tested. We also thank all contributors to the EMPIAR data repository.

## 8 Supplementary Materials

### 8.1 Particle Extraction Box Sizes

Across our test set, each dataset had a unique pixel size. Once particles had been picked and pre-filtered, we extracted them at a box size that would match as close as possible to the physical box extent of the input 3D reference map. For example, the active GPCR reference maps have a box size of 320 pixels and a pixel size of 1.034 Å, yielding a box extent of 330.9Å. Therefore, a dataset with a pixel size of 0.83Å would require particles to be extracted at a box size of 376 to match. Note that in practice, if all datasets are collected at the same pixel size, they can all be extracted at the same box size as well. For inactive GPCR datasets: Particles were extracted with a box extent of 222.1Å. For this class, a smaller box size resulted in cleaner reconstructions as a larger box size included neighboring particles.

### 8.2 Generation of Low-Resolution Reference Density Maps and Local Refinement Masks

Low resolution 3D reference maps were created from a manual processing of a single exemplar dataset per class. For the active Nb35 class, the reference was generated using the final Non-Uniform Refinement from manual processing of EMPIAR-10346. A generous mask was used to remove the flexible and disordered region of Nb35 that appeared to be present in reconstructions of this target (and others utilizing Nb35). The resulting volume was then lowpass filtered to 15Å. For the active scFv class, the reference was generated using the final Non-Uniform Refinement from manual processing of EMPIAR-11119. This volume was lowpass filtered to 15Å. For the active no-Fab class, the active scFv reference was masked to remove the scFv. For the inactive GPCR class, the final Non-Uniform Refinement from manual processing of EMPIAR-11134 was lowpass filtered to 15Å.

To generate masks for local refinements, the same exemplar datasets were used. The final NonUniform Refinements from EMPIAR-10346 (active Nb35), EMPIAR-11119 (active scFv and active no-Fab), and EMPIAR-11134 (inactive) were segmented in ChimeraX to isolate only the receptor density, imported into CryoSPARC, lowpass filtered to 15Å, thresholded, dilated (3 px) and softened (15 px).

### 8.3 Generation of Junk Volumes for Decoy Classification

Junk volumes are used during decoy classification. We generated four junk volumes, and used the same four volumes for all active and inactive target classes. Two of these volumes, representative of an empty micelle and G-protein complex, were generated by subtracting out various portions of a Non-Uniform Refinement map from EMPIAR-10288, lowpass filtered to 15Å. For the other two volumes, we took a large subset of junk particle picks (on ice/ethane, background, etc.) that were rejected during manual processing of EMPIAR-10288 and ran an Ab-Initio Reconstruction job, terminating it after only a few iterations and keeping the output volumes.

### 8.4 Generation of Masks for Computing FSC Curves

To generate masks for computing global and local gold standard FSC curves, the final Non-Uniform Refinement and Local Refinement volumes for each dataset were segmented in ChimeraX to remove micelle density. These volumes were then imported into CryoSPARC, lowpass filtered to 15Å, and converted from a map to a mask at an appropriate threshold as determined by manual adjustment in ChimeraX. The threshold was chosen to ensure that after a soft edge (15 px) and dilation (3 px) were applied, the GPCR complex or receptor density would be fully encapsulated. A single mask was created for each dataset, and this mask was used to compute FSC curves and FSC resolution estimates for the deposited maps (where half-maps were available) and for results from our automated processing for that dataset, to ensure that resolution comparison remained fair.

### 8.5 Computational Hardware

All datasets were processed on a node with an AMD Threadripper Pro 5975WX CPU, 2× NVIDIA RTX 4090 GPUs, and 256 GB DDR4 RAM. Two 4 TB NVMe SSDs in RAID 0 were utilized for CryoSPARC’s internal particle caching system, while raw movies and motion corrected micrographs were stored on a large ZFS HDD array accessed via 10 Gb Ethernet. The preprocessing and Reference Based Motion Correction stages for five datasets were re-processed on a node with AMD EPYC 7F53 CPU, 8x NVIDIA A100 GPUs, 512GB DDR4 RAM, and 2x 8 TB NVMe SSDs in RAID 0.

### 8.6 Supplementary Tables

The following tables provide additional data regarding test datasets, results, and comparisons.

---

**Table S1.**
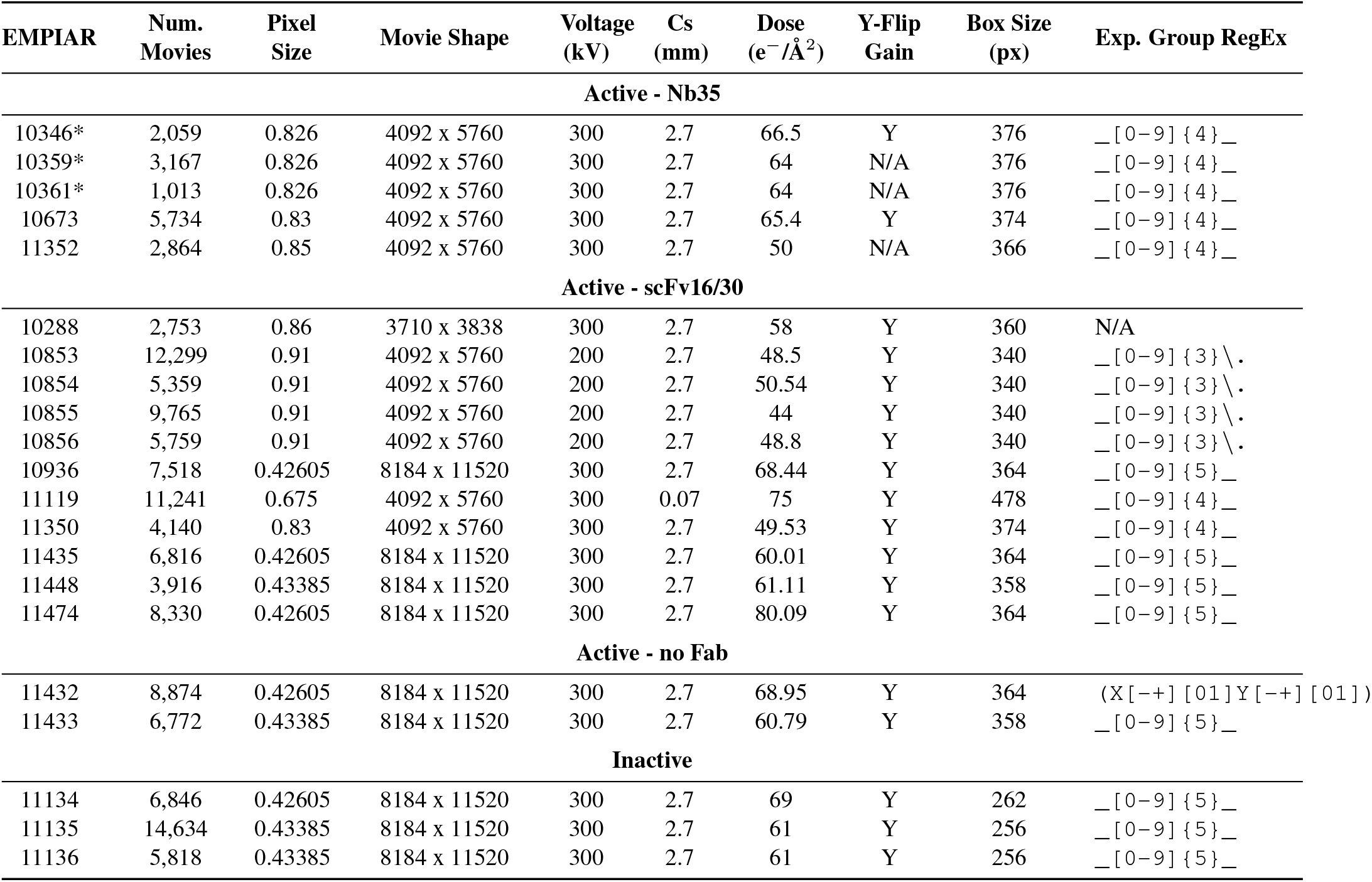
Microscope imaging parameters for each EMPIAR dataset in the test set. Asterisk indicates datasets that include phase plate data in the deposition; our analysis only uses non phase plate data. The final column is the regular expression used to split images into exposure groups based on file name.

**Table S2.** Full automated processing results across 21 GPCR datasets. Our automated workflow achieves equal or better resolution in 17 out of the 21 datasets with six improvements of 0.3Å or greater. FSC and cFAR were measured using a single soft mask for each dataset, so that the values are directly comparable (see Section 8.4). Asterisks indicate datasets for which half-maps were not deposited to EMDB, so the deposition-reported resolution is used instead. The fourth column provides resolution values from our automated processing before Reference Based Motion Correction (RBMC), showing that this step produces improvements for only a few datasets and could be omitted. The final column indicates which CTF abberations (3rd order: tilt, trefoil; 4th order: tetrafoil, spherical) were corrected by the automated workflow.

| EMPIAR | GSFSC Resolution (Å) |  |  |  | cFAR Score |  |  | CTF aberr.<br>corrected |
| --- | --- | --- | --- | --- | --- | --- | --- | --- |
|  | Deposited | Auto.<br>Global | Auto. w/o<br>RBMC | Auto.<br>Local | Deposited | Auto.<br>Global | Auto.<br>Local |  |
| Active - Nb35 |  |  |  |  |  |  |  |  |
| 10346 | 3.1 | 3.0 ↑ | 3.0 | 3.2 | 0.53 | 0.63 ↑ | 0.51 | 3rd |
| 10359 | 2.7 | 2.6 ↑ | 2.6 | 2.6 | 0.55 | 0.57 ↑ | 0.53 | 3rd |
| 10361 | 2.8 | 2.9 ↓ | 2.9 | 3.0 | 0.58 | 0.58 ≈ | 0.53 | 3rd, 4th |
| 10673 | 2.2 | 2.1 ↑ | 2.2 | 2.4 | 0.52 | 0.56 ↑ | 0.50 | 3rd |
| 11352 | 3.1 | 2.8 ↑ | 3.0 | 3.0 | 0.42 | 0.65 ↑ | 0.61 | 3rd |
| Active - scFv16/30 |  |  |  |  |  |  |  |  |
| 10288* | 3.0 | 2.7 ↑ | 2.7 | 3.0 | — | 0.60 | 0.40 | 3rd, 4th |
| 10853 | 2.5 | 2.6 ↓ | 2.7 | 2.8 | 0.41 | 0.65 ↑ | 0.40 | 3rd, 4th |
| 10854 | 3.0 | 2.9 ↑ | 3.0 | 3.2 | 0.65 | 0.75 ↑ | 0.70 | 3rd, 4th |
| 10855 | 2.6 | 2.6 ≈ | 2.7 | 2.8 | 0.56 | 0.65 ↑ | 0.50 | 3rd, 4th |
| 10856 | 2.6 | 2.6 ≈ | 2.7 | 2.8 | 0.71 | 0.82 ↑ | 0.67 | 3rd, 4th |
| 10936* | 2.9 | 2.8 ↑ | 2.8 | 3.0 | — | 0.41 | 0.25 | 3rd, 4th |
| 11119 | 3.3 | 3.0 ↑ | 3.0 | 3.4 | 0.71 | 0.72 ↑ | 0.49 | 3rd |
| 11350 | 3.5 | 2.8 ↑ | 2.8 | 3.2 | 0.25 | 0.66 ↑ | 0.47 | 3rd |
| 11435* | 3.5 | 2.9 ↑ | 2.9 | 3.1 | — | 0.78 | 0.64 | 3rd, 4th |
| 11448* | 2.9 | 2.9 ≈ | 3.0 | 3.0 | — | 0.50 | 0.29 | 3rd |
| 11474 | 3.0 | 2.6 ↑ | 2.6 | 2.9 | 0.57 | 0.62 ↑ | 0.44 | 3rd, 4th |
| Active - no Fab |  |  |  |  |  |  |  |  |
| 11432* | 3.0 | 2.9 ↑ | 2.9 | 2.7 | — | 0.35 | 0.41 | 3rd, 4th |
| 11433* | 2.7 | 2.9 ↓ | 2.9 | 2.9 | — | 0.76 | 0.65 | 3rd, 4th |
| Inactive |  |  |  |  |  |  |  |  |
| 11134 | 3.1 | 3.1 ≈ | 3.1 | 3.1 | 0.60 | 0.52 ↓ | 0.63 | 3rd, 4th |
| 11135 | 2.8 | 2.8 ≈ | 2.8 | 2.7 | 0.15 | 0.13 ↓ | 0.22 | 3rd, 4th |
| 11136 | 2.5 | 2.6 ↓ | 2.6 | 2.5 | 0.41 | 0.51 ↑ | 0.54 | 3rd, 4th |

**Table S3.** Particle counts (in millions) remaining after each stage of processing, across the 21 EMPIAR datasets in the test set. The automated workflow is able to successfully curate datasets ranging from 1 to 18 million initial particles, while the fraction of final particles retained ranges from 2 to 28 percent.

| EMPIAR<br>Entry | Particles Remaining After Stage (Millions) |  |  |  |  | Final Percent<br>Retained |
| --- | --- | --- | --- | --- | --- | --- |
|  | Template<br>picking | Junk<br>detector | Auto<br>inspect | Decoy clas-<br>sification | Final<br>stack |  |
| Active - Nb35 |  |  |  |  |  |  |
| 10346 | 1.61 | 1.54 | 0.71 | 0.22 | 0.14 | 9% |
| 10359 | 3.42 | 3.29 | 2.17 | 0.89 | 0.77 | 23% |
| 10361 | 0.96 | 0.93 | 0.66 | 0.27 | 0.21 | 21% |
| 10673 | 4.18 | 4.12 | 3.00 | 1.18 | 0.58 | 14% |
| 11352 | 2.20 | 1.90 | 0.88 | 0.27 | 0.15 | 7% |
| Active - scFv16/30 |  |  |  |  |  |  |
| 10288 | 1.55 | 1.52 | 0.95 | 0.57 | 0.44 | 28% |
| 10853 | 11.8 | 11.3 | 9.40 | 3.79 | 3.10 | 26% |
| 10854 | 5.06 | 4.69 | 2.20 | 1.02 | 0.85 | 17% |
| 10855 | 10.0 | 9.77 | 8.18 | 3.59 | 2.40 | 24% |
| 10856 | 6.20 | 6.21 | 5.13 | 2.11 | 1.50 | 24% |
| 10936 | 6.42 | 6.08 | 3.04 | 1.24 | 0.67 | 10% |
| 11119 | 5.31 | 5.00 | 2.41 | 0.95 | 0.67 | 13% |
| 11350 | 3.55 | 3.48 | 1.77 | 0.46 | 0.26 | 7% |
| 11435 | 5.52 | 5.00 | 2.17 | 0.63 | 0.40 | 7% |
| 11448 | 3.56 | 3.51 | 2.42 | 1.01 | 0.87 | 24% |
| 11474 | 6.26 | 6.09 | 3.97 | 1.90 | 1.58 | 25% |
| Active - no Fab |  |  |  |  |  |  |
| 11432 | 4.27 | 4.10 | 2.74 | 1.14 | 0.70 | 16% |
| 11433 | 7.01 | 6.94 | 5.78 | 2.23 | 1.60 | 23% |
| Inactive |  |  |  |  |  |  |
| 11134 | 10.80 | 10.68 | 7.30 | 1.22 | 0.96 | 9% |
| 11135 | 18.05 | 17.44 | 13.04 | 0.66 | 0.43 | 2% |
| 11136 | 9.10 | 9.02 | 6.09 | 1.03 | 0.80 | 9% |

**Table S4.** Processing times in hours for each test dataset, using a 2-GPU workstation. Times are shown for each stage of processing, as well as the total run time. For five datasets, times are also shown for preprocessing and RBMC stages using an 8-GPU server (grey), showing that 2-3x speedup in these stages is possible and an overall speedup of up to 1.7x by using additional hardware.

| EMPIAR Entry | Preprocessing | Picking | Particle Curation | Final Re-finements | RBMC | Total | # of movies | Movie shape |
| --- | --- | --- | --- | --- | --- | --- | --- | --- |
| **Active - Nb35** |  |  |  |  |  |  |  |  |
| 10346 | 3.7 | 0.6 | 3.3 | 1.0 | 2.7 | 11.3 | 2,059 | 4092 x 5760 |
| 10359 | 6.2 | 1.0 | 5.3 | 3.2 | 5.8 | 21.5 | 3,167 | 4092 x 5760 |
| 10361 | 1.8 | 0.6 | 3.6 | 1.2 | 2.2 | 9.4 | 1,013 | 4092 x 5760 |
| (8-GPU) | 0.7 |  |  |  | 1.5 | 7.6 |  |  |
| 10673 | 9.8 | 0.9 | 6.3 | 2.6 | 6.3 | 25.9 | 5,734 | 4092 x 5760 |
| 11352 | 4.2 | 0.5 | 3.6 | 1.1 | 1.9 | 11.3 | 2,864 | 4092 x 5760 |
| **Active - scFv16/30** |  |  |  |  |  |  |  |  |
| 10288 | 2.5 | 0.7 | 3.7 | 1.7 | 2.3 | 10.9 | 2,753 | 3710 x 3838 |
| 10853 | 17.9 | 2.5 | 13.0 | 9.7 | 16.5 | 59.6 | 12,299 | 4092 x 5760 |
| (8-GPU) | 6.7 |  |  |  | 9.0 | 40.9 |  |  |
| 10854 | 8.0 | 1.4 | 6.0 | 2.6 | 4.6 | 22.6 | 5,359 | 4092 x 5760 |
| 10855 | 14.3 | 2.0 | 11.1 | 7.5 | 14.3 | 49.2 | 9,765 | 4092 x 5760 |
| (8-GPU) | 5.4 |  |  |  | 7.7 | 33.7 |  |  |
| 10856 | 8.3 | 1.7 | 8.1 | 3.6 | 8.3 | 30.0 | 5,759 | 4092 x 5760 |
| 10936 | 17.0 | 1.5 | 6.3 | 2.3 | 11.1 | 38.2 | 7,518 | 8184 x 11520 |
| (8-GPU) | 6.4 |  |  |  | 8.6 | 25.1 |  |  |
| 11119 | 16.1 | 1.7 | 5.7 | 2.3 | 7.3 | 33.1 | 11,241 | 4092 x 5760 |
| 11350 | 5.5 | 0.9 | 4.9 | 1.3 | 2.5 | 15.1 | 4,140 | 4092 x 5760 |
| 11435 | 14.8 | 1.1 | 5.0 | 1.6 | 7.8 | 30.3 | 6,816 | 8184 x 11520 |
| 11448 | 9.1 | 1.1 | 5.8 | 2.0 | 18.3 | 36.3 | 3,916 | 8184 x 11520 |
| 11474 | 19.7 | 1.2 | 6.6 | 5.7 | 30.6 | 63.8 | 8,330 | 8184 x 11520 |
| **Active - no Fab** |  |  |  |  |  |  |  |  |
| 11432 | 21.8 | 1.0 | 6.2 | 2.5 | 10.8 | 42.3 | 8,874 | 8184 x 11520 |
| 11433 | 15.4 | 1.6 | 9.5 | 4.3 | 33.0 | 63.8 | 6,772 | 8184 x 11520 |
| **Inactive** |  |  |  |  |  |  |  |  |
| 11134 | 15.6 | 1.4 | 10.6 | 2.6 | 9.0 | 39.2 | 6,846 | 8184 x 11520 |
| 11135 | 35.9 | 2.2 | 13.3 | 1.2 | 13.1 | 65.7 | 14,634 | 8184 x 11520 |
| (8-GPU) | 13.9 |  |  |  | 9.3 | 39.9 |  |  |
| 11136 | 14.7 | 1.3 | 9.4 | 1.6 | 9.6 | 36.6 | 5,818 | 8184 x 11520 |

**Table S5.** Amount of storage space in TB used for processing each test dataset. The second column shows the size of the raw data, and the third column shows the size of all output created by CryoSPARC (i.e. project data size) during processing. At most, 11 TB are needed to store the raw plus project data for EMPIAR-11135, with the majority of datasets requiring substantially less space.

| Dataset | Size of<br>EMPIAR<br>Deposition (TB) | CryoSPARC<br>Project<br>Data (TB) | Total<br>Amount<br>of Storage<br>Used (TB) | Num.<br>Movies | Movie Shape | Initial<br>Particle<br>Picks | Final<br>Particle<br>Picks |
| --- | --- | --- | --- | --- | --- | --- | --- |
| **Active - Nb35** |  |  |  |  |  |  |  |
| 10346 | 1.4 | 0.4 | 1.7 | 2,059 | 4092 x 5760 | 1.61M | 0.13M |
| 10359 | 2.0 | 0.8 | 2.8 | 3,167 | 4092 x 5760 | 3.42M | 0.76M |
| 10361 | 0.6 | 0.3 | 0.9 | 1,013 | 4092 x 5760 | 0.96M | 0.21M |
| 10673 | 1.9 | 1.1 | 3.0 | 5,734 | 4092 x 5760 | 4.18M | 0.58M |
| 11352 | 0.7 | 0.5 | 1.1 | 2,864 | 4092 x 5760 | 2.20M | 0.15M |
| **Active - scFv16/30** |  |  |  |  |  |  |  |
| 10288 | 0.5 | 0.5 | 0.9 | 2,753 | 3710 x 3838 | 1.55M | 0.44M |
| 10853 | 3.6 | 3.0 | 6.6 | 12,299 | 4092 x 5760 | 11.80M | 3.10M |
| 10854 | 1.6 | 1.1 | 2.7 | 5,359 | 4092 x 5760 | 5.06M | 0.85M |
| 10855 | 2.8 | 2.5 | 5.3 | 9,765 | 4092 x 5760 | 10.00M | 2.40M |
| 10856 | 1.7 | 1.5 | 3.2 | 5,759 | 4092 x 5760 | 6.20M | 1.50M |
| 10936 | 4.1 | 1.3 | 5.4 | 7,518 | 8184 x 11520 | 6.42M | 0.67M |
| 11119 | 3.1 | 1.6 | 4.7 | 11,241 | 4092 x 5760 | 5.31M | 0.67M |
| 11350 | 1.4 | 0.7 | 2.1 | 4,140 | 4092 x 5760 | 3.55M | 0.26M |
| 11435 | 3.4 | 1.0 | 4.4 | 6,816 | 8184 x 11520 | 5.52M | 0.40M |
| 11448 | 2.0 | 0.9 | 2.9 | 3,916 | 8184 x 11520 | 3.56M | 0.87M |
| 11474 | 5.2 | 1.7 | 6.9 | 8,330 | 8184 x 11520 | 6.26M | 1.58M |
| **Active - no Fab** |  |  |  |  |  |  |  |
| 11432 | 5.3 | 1.4 | 6.7 | 8,874 | 8184 x 11520 | 4.27M | 0.70M |
| 11433 | 3.6 | 1.7 | 5.3 | 6,772 | 8184 x 11520 | 7.01M | 1.06M |
| **Inactive** |  |  |  |  |  |  |  |
| 11134 | 3.8 | 1.7 | 5.5 | 6,846 | 8184 x 11520 | 10.80M | 0.96M |
| 11135 | 8.1 | 2.4 | 10.5 | 14,634 | 8184 x 11520 | 18.05M | 0.43M |
| 11136 | 3.3 | 1.2 | 4.5 | 5,818 | 8184 x 11520 | 9.10M | 0.80M |

**Table S6.** Manual processing workflows that were used by the original authors for each EMPIAR dataset. Superscripts indicate software: ^1^CryoSPARC, ^2^RELION, ^3^cisTEM.

| EMPIAR | Software(s) | Preprocessing | Picking | Particle Curation | Final Refinements | Polishing |
| --- | --- | --- | --- | --- | --- | --- |
| **Active - Nb35** |  |  |  |  |  |  |
| 10346 | CryoSPARC, RELION | MotionCor2, GCTF | LoG<sup>2</sup> | 2D-class<sup>2</sup>, ab-initio<sup>1</sup>, 3D-class<sup>2</sup> | Micelle subtraction<sup>2</sup>, 3D-refine<sup>2</sup> | Y |
| 10359 | CryoSPARC, RELION | MotionCor2, GCTF | crYOLO | 2D-class<sup>2</sup>, ab-initio<sup>1</sup>, 3D-class<sup>2</sup>, 3D-class (no align)<sup>2</sup> | 3D-refine<sup>2</sup> | Y |
| 10361 | RELION | MotionCor2, GCTF | LoG + Template | 3D-class | 3D-refine | Y |
| 10673 | RELION | MotionCor2, GCTF | LoG | 2D-class, ab-initio, 3D-class | 3D-refine, CTF refine, local ref | Y |
| 11352 | RELION | MotionCor2, CTFFind4 | Manual, template | 2D-class, 3D-class, 3D-class (no align) | 3D-refine, micelle subtraction, CTF refine | Y |
| **Active - scFv16/30** |  |  |  |  |  |  |
| 10288 | cisTEM, RELION | MotionCor2, CTFFind4 | LoG<sup>2</sup> | 2D-class<sup>2</sup>, 3D-class w/ LPF EMD-7868<sup>2</sup> | 3D-refine<sup>3</sup> | N |
| 10853 | CryoSPARC | Patch Motion, Patch CTF | Blob/template, TOPAZ | 2D-class, ab-initio, hetref | NU-ref, CTF refine, local ref | N |
| 10854 | CryoSPARC | Patch Motion, Patch CTF | Blob/template, TOPAZ | 2D-class, ab-initio, hetref | NU-ref, CTF refine, local ref | N |
| 10855 | CryoSPARC | Patch Motion, Patch CTF | Blob/template, TOPAZ | 2D-class, ab-initio, hetref | NU-ref, CTF refine, local ref | N |
| 10856 | CryoSPARC | Patch Motion, Patch CTF | Blob/template, TOPAZ | 2D-class, ab-initio, hetref | NU-ref, CTF refine, local ref | N |
| 10936 | CryoSPARC, RELION | MotionCor2, CTFFind4 | LoG<sup>2</sup> | 2D-class<sup>1</sup>, ab-initio<sup>1</sup>, hetref<sup>1</sup> | NU-ref, CTF refine<sup>1</sup>, local ref<sup>1</sup> | Y |
| 11119 | RELION | MotionCor2, GCTF | LoG, Template | 2D-class, 3D-class, 3D-class (no align) | 3D-refine, SIDESPLITTER, local ref | Y |
| 11350 | RELION | RELION MotCor, CTFFind4 | — | 2D-class, 3D-class, 3D-class (no align) | 3D-refine, micelle subtraction, CTF refine, local ref | Y |
| 11435 | RELION | MotionCor2, CTFFind4 | Template | 2D-class, 3D-class | 3D-refine, CTF refine | Y |
| 11448 | CryoSPARC | Patch Motion, Patch CTF | — | 2D-class, ab-initio, hetref | NU-ref, CTF refine, local ref | N |
| 11474 | CryoSPARC, RELION | MotionCor2, CTFFind4 | — | 2D-class<sup>1</sup>, ab-initio<sup>1</sup>, hetref<sup>1</sup> | NU-ref, CTF refine<sup>1</sup>, local ref | Y |
| **Active - no Fab** |  |  |  |  |  |  |
| 11432 | CryoSPARC | Patch Motion, Patch CTF | — | 2D-class, ab-initio, hetref | NU-ref, CTF refine, local ref | N |
| 11433 | CryoSPARC | Patch Motion, Patch CTF | — | 2D-class, ab-initio, hetref | NU-ref, CTF refine, local ref | N |
| **Inactive** |  |  |  |  |  |  |
| 11134 | CryoSPARC, RELION | MotionCor2, CTFFind4 | LoG | 2D-class<sup>1</sup>, ab-initio<sup>1</sup>, hetref<sup>1</sup> | NU-ref, CTF refine<sup>1</sup>, local ref | Y |
| 11135 | CryoSPARC, RELION | MotionCor2, CTFFind4 | LoG | 2D-class<sup>1</sup>, ab-initio<sup>1</sup>, hetref<sup>1</sup> | NU-ref, CTF refine<sup>1</sup>, local ref | Y |
| 11136 | CryoSPARC, RELION | MotionCor2, CTFFind4 | LoG | 2D-class<sup>1</sup>, ab-initio<sup>1</sup>, hetref<sup>1</sup> | NU-ref, CTF refine<sup>1</sup>, local ref | Y |

**Table S7.**
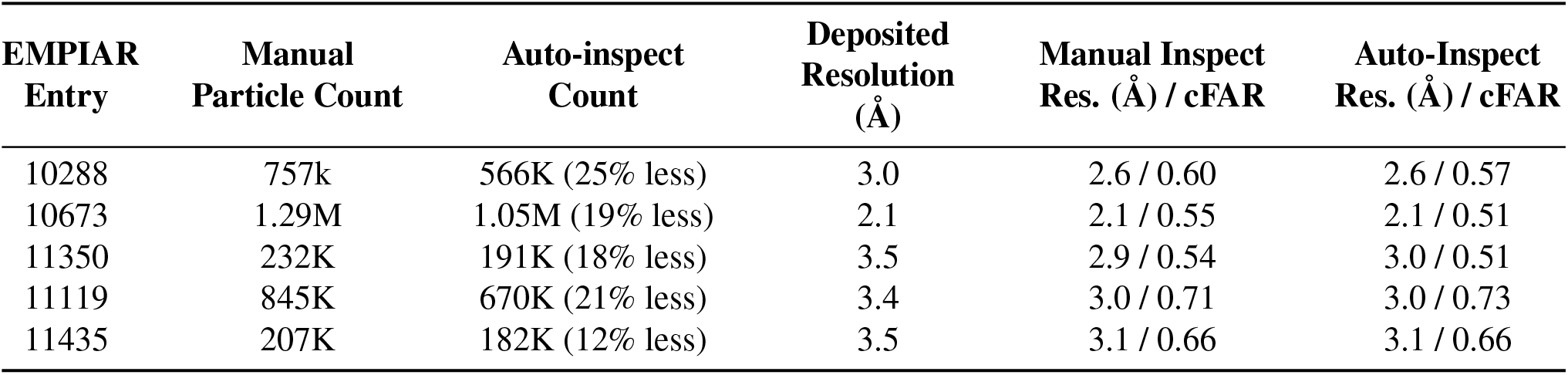
Comparison of manual particle inspection and auto cluster mode in Inspect Particle Picks (Section 7.2.3) across five test datasets. The second column shows the number of particles retained after manual particle inspection and setting of pick score thresholds while the third column shows the number retained by auto cluster mode. Resolutions and cFAR scores from auto clustering are similar to manual inspection.

**Table S8.** Comparison of template picking on denoised micrographs (third column) against Topaz [46], using a general GPCR model trained on additional datasets (fourth column) or a dataset-specific model trained on each test dataset (fifth column). Results achieved by template picking are similar or improved relative to Topaz, despite the simplicity of the template picking.

| EMPIAR Entry | Deposited Resolution (Å) / cFAR | Template Picking Resolution (Å) / cFAR | TOPAZ General Model Resolution (Å) / cFAR | TOPAZ Dataset Specific Model Resolution (Å) / cFAR |
| --- | --- | --- | --- | --- |
| 10359 | 2.7 / 0.55 | 2.5 / 0.59 | 2.6 / 0.52 | 2.6 / 0.54 |
| 10361 | 2.8 / 0.58 | 2.8 / 0.58 | 2.9 / 0.51 | 3.0 / 0.54 |
| 10673 | 2.2 / 0.52 | 2.1 / 0.59 | 2.1 / 0.60 | 2.2 / 0.62 |
| 10853 | 2.5 / 0.41 | 2.7 / 0.59 | 2.7 / 0.59 | 2.7 / 0.53 |
| 10936 | 2.9 / - | 2.8 / 0.34 | 2.7 / 0.36 | 2.8 / 0.39 |

## References

[1] E. Y. D. Chua, J. H. Mendez, M. Rapp, S. L. Ilca, Y. Z. Tan, K. Maruthi, H. Kuang, C. M. Zimanyi, A. Cheng, E. T. Eng, A. J. Noble, C. S. Potter, and B. Carragher, “Better, Faster, Cheaper: Recent Advances in Cryo-Electron Microscopy.,” Annual review of biochemistry, vol. 91, pp. 1–32, June 2022. 1

[2] V. I. Cushing, A. F. Koh, J. Feng, K. Jurgaityte, A. Bondke, S. H. B. Kroll, M. Barbazanges, B. Scheiper, A. K. Bahl, A. G. M. Barrett, S. Ali, A. Kotecha, and B. J. Greber, “High-resolution cryo-EM of the human CDK-activating kinase for structure-based drug design,” Nature Communications, vol. 15, p. 2265, Mar. 2024. Publisher: Nature Publishing Group. 1

[3] S. Bangaru, A. Antanasijevic, N. Kose, L. M. Sewall, A. M. Jackson, N. Suryadevara, X. Zhan, J. L. Torres, J. Copps, A. T. de la Peña, J. E. Crowe, and A. B. Ward, “Structural mapping of antibody landscapes to human betacoronavirus spike proteins,” Science Advances, vol. 8, p. eabn2911, May 2022. Publisher: American Association for the Advancement of Science. 1

[4] M. M. Papasergi-Scott, G. Pérez-Hernández, H. Batebi, Y. Gao, G. Eskici, A. B. Seven, O. Panova, D. Hilger, M. Casiraghi, F. He, L. Maul, P. Gmeiner, B. K. Kobilka, P. W. Hildebrand, and G. Skiniotis, “Time-resolved cryo-EM of G-protein activation by a GPCR,” Nature, vol. 629, pp. 1182–1191, May 2024. Publisher: Nature Publishing Group. 1

[5] I. D. Lutz, S. Wang, C. Norn, A. Courbet, A. J. Borst, Y. T. Zhao, A. Dosey, L. Cao, J. Xu, E. M. Leaf, C. Treichel, P. Litvicov, Z. Li, A. D. Goodson, P. Rivera-Sánchez, A.-M. Bratovianu, M. Baek, N. P. King, H. Ruohola-Baker, and D. Baker, “Top-down design of protein architectures with reinforcement learning,” Science, vol. 380, pp. 266–273, Apr. 2023. Publisher: American Association for the Advancement of Science. 1

[6] “CryoSPARC: Live.” https://cryosparc.com/live. 2, 17

[7] D. Tegunov and P. Cramer, “Real-time cryo-electron microscopy data preprocessing with Warp,” Nature Methods, vol. 16, pp. 1146– 1152, Nov. 2019. 2, 22, 24

[8] M. Stabrin, F. Schoenfeld, T. Wagner, S. Pospich, C. Gatsogiannis, and S. Raunser, “TranSPHIRE: automated and feedback-optimized on-the-fly processing for cryo-EM,” Nature Communications, vol. 11, p. 5716, Nov. 2020. Publisher: Nature Publishing Group. 2, 26

[9] J. Gómez-Blanco, J. M. de la Rosa-Trevín, R. Marabini, L. del Cano, A. Jiménez, M. Martínez, R. Melero, T. Majtner, D. Maluenda, J. Mota, Y. Rancel, E. Ramírez-Aportela, J. L. Vilas, M. Carroni, S. Fleischmann, E. Lindahl, A. W. Ashton, M. Basham, D. K. Clare, K. Savage, C. A. Siebert, G. G. Sharov, C. O. S. Sorzano, P. Conesa, and J. M. Carazo, “Using Scipion for stream image processing at Cryo-EM facilities,” Journal of Structural Biology, vol. 204, pp. 457–463, Dec. 2018. 2

[10] “Getting Started — cryosparc-tools.” https://tools.cryosparc.com/intro.html. 2

[11] J. Zivanov, T. Nakane, B. Forsberg, D. Kimanius, W. Hagen, and E. Lindahl, “New tools for automated high-resolution cryo-em structure determination in RELION-3,” eLife, vol. 7, e42166, 2018. 2

[12] J. M. de la Rosa-Trevín, A. Quintana, L. del Cano, A. Zaldívar, I. Foche, J. Gutiérrez, J. Gómez-Blanco, J. Burguet-Castell, J. Cuenca-Alba, V. Abrishami, J. Vargas, J. Otón, G. Sharov, J. L. Vilas, J. Navas, P. Conesa, M. Kazemi, R. Marabini, C. O. S. Sorzano, and J. M. Carazo, “Scipion: A software framework toward integration, reproducibility and validation in 3D electron microscopy,” Journal of Structural Biology, vol. 195, pp. 93–99, July 2016. 2

[13] D. Kimanius, L. Dong, G. Sharov, T. Nakane, and S. H. W. Scheres, “New tools for automated cryo-EM single-particle analysis in RELION-4.0,” Biochemical Journal, vol. 478, pp. 4169–4185, Dec. 2021. 2, 26

[14] Y. Cheng, X. Huang, B. Xu, and W. Ding, “AutoEMage: automatic data transfer, preprocessing, real-time display and monitoring in cryo-EM,” Journal of Applied Crystallography, vol. 56, pp. 1865–1873, Dec. 2023. Publisher: International Union of Crystallography. 2

[15] Y. Yan, S. Fan, F. Yuan, and H. Shen, “A comprehensive foundation model for cryo-EM image processing,” Nov. 2024. Pages: 2024.11.04.621604 Section: New Results. 2

[16] Y. Li, J. N. Cash, J. J. G. Tesmer, and M. A. Cianfrocco, “High-Throughput Cryo-EM Enabled by User-Free Preprocessing Routines,” Structure, vol. 28, pp. 858–869.e3, July 2020. 2, 26

[17] J.-H. Schäfer, A. Calza, K. Hom, P. Damodar, R. Peng, N. Bogdanović, G. C. Lander, S. M. Stagg, and M. A. Cianfrocco, “CryoSift – An accessible and automated CNN-driven tool for cryo-EM 2D class selection,” bioRxiv, p. 2025.07.28.667259, Sept. 2025. 2, 26

[18] M. Zhang, T. Chen, X. Lu, X. Lan, Z. Chen, and S. Lu, “G protein-coupled receptors (GPCRs): advances in structures, mechanisms and drug discovery,” Signal Transduction and Targeted Therapy, vol. 9, p. 88, Apr. 2024. Publisher: Nature Publishing Group. 3

[19] M. Congreve, C. d. Graaf, N. A. Swain, and C. G. Tate, “Impact of GPCR Structures on Drug Discovery,” Cell, vol. 181, pp. 81–91, Apr. 2020. Publisher: Elsevier. 3

[20] J. Caroli, S. N. Andreassen, J. S. Lorente, B. Xiao, G. Pándy-Szekeres, and D. E. Gloriam, “An online GPCR drug discovery resource,” npj Drug Discovery, vol. 2, p. 17, July 2025. Publisher: Nature Publishing Group. 3

[21] K. Zhang, H. Wu, N. Hoppe, A. Manglik, and Y. Cheng, “Fusion protein strategies for cryo-EM study of G protein-coupled receptors,” Nature Communications, vol. 13, p. 4366, July 2022. Publisher: Nature Publishing Group. 3

[22] M. J. Robertson, M. M. Papasergi-Scott, F. He, A. B. Seven, J. G. Meyerowitz, O. Panova, M. C. Peroto, T. Che, and G. Skiniotis, “Structure determination of inactive-state GPCRs with a universal nanobody,” Nature Structural & Molecular Biology, vol. 29, pp. 1188–1195, Dec. 2022. Publisher: Nature Publishing Group. 3

[23] A. Iudin, P. K. Korir, S. Somasundharam, S. Weyand, C. Cattavitello, N. Fonseca, O. Salih, G. Kleywegt, and A. Patwardhan, “EMPIAR: the Electron Microscopy Public Image Archive,” Nucleic Acids Research, vol. 51, pp. D1503–D1511, Jan. 2023. 3, 34

[24] H. M. Berman, J. Westbrook, Z. Feng, G. Gilliland, T. N. Bhat, H. Weissig, I. N. Shindyalov, and P. E. Bourne, “The Protein Data Bank,” Nucleic Acids Research, vol. 28, pp. 235–242, Jan. 2000. 4

[25] The wwPDB Consortium, “EMDB—the Electron Microscopy Data Bank,” Nucleic Acids Research, vol. 52, pp. D456–D465, Jan. 2024. 4, 8

[26] P. Zhao, Y.-L. Liang, M. J. Belousoff, G. Deganutti, M. M. Fletcher, F. S. Willard, M. G. Bell, M. E. Christe, K. W. Sloop, A. Inoue, T. T. Truong, L. Clydesdale, S. G. B. Furness, A. Christopoulos, M.-W. Wang, L. J. Miller, C. A. Reynolds, R. Danev, P. M. Sexton, and D. Wootten, “Activation of the glp-1 receptor by a non-peptidic agonist,” Nature, vol. 577, p. 432–436, Jan. 2020. 4

[27] Y.-L. Liang, M. J. Belousoff, P. Zhao, C. Koole, M. M. Fletcher, T. T. Truong, V. Julita, G. Christopoulos, H. E. Xu, Y. Zhang, M. Khoshouei, A. Christopoulos, R. Danev, P. M. Sexton, and D. Wootten, “Toward a structural understanding of class b gpcr peptide binding and activation,” Molecular Cell, vol. 77, pp. 656–668.e5, Feb. 2020. 4

[28] X. Zhang, M. J. Belousoff, P. Zhao, A. J. Kooistra, T. T. Truong, S. Y. Ang, C. R. Underwood, T. Egebjerg, P. Šenel, G. D. Stewart, Y.-L. Liang, A. Glukhova, H. Venugopal, A. Christopoulos, S. G. Furness, L. J. Miller, S. Reedtz-Runge, C. J. Langmead, D. E. Gloriam, R. Danev, P. M. Sexton, and D. Wootten, “Differential glp-1r binding and activation by peptide and non-peptide agonists,” Molecular Cell, vol. 80, pp. 485–500.e7, Nov. 2020. 4

[29] C. Nagiri, K. Kobayashi, A. Tomita, M. Kato, K. Kobayashi, K. Yamashita, T. Nishizawa, A. Inoue, W. Shihoya, and O. Nureki, “Cryo-em structure of the 3-adrenergic receptor reveals the molecular basis of subtype selectivity,” Molecular Cell, vol. 81, pp. 3205–3215.e5, Aug. 2021. 4

[30] K. Krishna Kumar, M. Shalev-Benami, M. J. Robertson, H. Hu, S. D. Banister, S. A. Hollingsworth, N. R. Latorraca, H. E. Kato, D. Hilger, S. Maeda, W. I. Weis, D. L. Farrens, R. O. Dror, S. V. Malhotra, B. K. Kobilka, and G. Skiniotis, “Structure of a signaling cannabinoid receptor 1-g protein complex,” Cell, vol. 176, pp. 448–458.e12, Jan. 2019. 4

[31] C. Cao, H. J. Kang, I. Singh, H. Chen, C. Zhang, W. Ye, B. W. Hayes, J. Liu, R. H. Gumpper, B. J. Bender, S. T. Slocum, B. E. Krumm, K. Lansu, J. D. McCorvy, W. K. Kroeze, J. G. English, J. F. DiBerto, R. H. J. Olsen, X.-P. Huang, S. Zhang, Y. Liu, K. Kim, J. Karpiak, L. Y. Jan, S. N. Abraham, J. Jin, B. K. Shoichet, J. F. Fay, and B. L. Roth, “Structure, function and pharmacology of human itch gpcrs,” Nature, vol. 600, p. 170–175, Nov. 2021. 4

[32] J. G. Meyerowitz, M. J. Robertson, X. Barros-Álvarez, O. Panova, R. M. Nwokonko, Y. Gao, and G. Skiniotis, “The oxytocin signaling complex reveals a molecular switch for cation dependence,” Nature Structural Molecular Biology, vol. 29, p. 274–281, Mar. 2022. 4

[33] R. Suno, Y. Sugita, K. Morimoto, H. Takazaki, H. Tsujimoto, M. Hirose, C. Suno-Ikeda, N. Nomura, T. Hino, A. Inoue, K. Iwasaki, T. Kato, S. Iwata, and T. Kobayashi, “Structural insights into the g protein selectivity revealed by the human ep3-gi signaling complex,” Cell Reports, vol. 40, p. 111323, Sept. 2022. 4, 14

[34] H. Akasaka, T. Tanaka, F. K. Sano, Y. Matsuzaki, W. Shihoya, and O. Nureki, “Structure of the active gi-coupled human lysophosphatidic acid receptor 1 complexed with a potent agonist,” Nature Communications, vol. 13, Sept. 2022. 4

[35] A. L. Kaplan, D. N. Confair, K. Kim, X. Barros-Álvarez, R. M. Rodriguiz, Y. Yang, O. S. Kweon, T. Che, J. D. McCorvy, D. N. Kamber, J. P. Phelan, L. C. Martins, V. M. Pogorelov, J. F. DiBerto, S. T. Slocum, X.-P. Huang, J. M. Kumar, M. J. Robertson, O. Panova, A. B. Seven, A. Q. Wetsel, W. C. Wetsel, J. J. Irwin, G. Skiniotis, B. K. Shoichet, B. L. Roth, and J. A. Ellman, “Bespoke library docking for 5-ht2a receptor agonists with antidepressant activity,” Nature, vol. 610, p. 582–591, Sept. 2022. 4

[36] C. Cao, X. Barros-Álvarez, S. Zhang, K. Kim, M. A. Dämgen, O. Panova, C.-M. Suomivuori, J. F. Fay, X. Zhong, B. E. Krumm, R. H. Gumpper, A. B. Seven, M. J. Robertson, N. J. Krogan, R. Hüttenhain, D. E. Nichols, R. O. Dror, G. Skiniotis, and B. L. Roth, “Signaling snapshots of a serotonin receptor activated by the prototypical psychedelic lsd,” Neuron, vol. 110, pp. 3154–3167.e7, Oct. 2022. 4

[37] K. Krishna Kumar, M. J. Robertson, E. Thadhani, H. Wang, C.-M. Suomivuori, A. S. Powers, L. Ji, S. P. Nikas, R. O. Dror, A. Inoue, A. Makriyannis, G. Skiniotis, and B. Kobilka, “Structural basis for activation of cb1 by an endocannabinoid analog,” Nature Communications, vol. 14, May 2023. 4

[38] X. Barros-Álvarez, R. M. Nwokonko, A. Vizurraga, D. Matzov, F. He, M. M. Papasergi-Scott, M. J. Robertson, O. Panova, E. H. Yardeni, A. B. Seven, F. E. Kwarcinski, H. Su, M. C. Peroto, J. G. Meyerowitz, M. Shalev-Benami, G. G. Tall, and G. Skiniotis, “The tethered peptide activation mechanism of adhesion gpcrs,” Nature, vol. 604, p. 757–762, Apr. 2022. 4

[39] M. J. Robertson, M. Papasergi-Scott, F. He, A. B. Seven, J. G. Meyerowitz, O. Panova, M. C. Peroto, T. Che, and G. Skiniotis, “Structure determination of inactive-state gpcrs with a universal nanobody,” Nov. 2021. 4

[40] “Tutorial: Orientation Diagnostics | CryoSPARC Guide.” https://guide.cryosparc.com/processing-data/tutorials-and-case-studies/tutorial-orientation-diagnostics, Nov. 2023. 8, 32

[41] E. C. Meng, T. D. Goddard, E. F. Pettersen, G. S. Couch, Z. J. Pearson, J. H. Morris, and T. E. Ferrin, “UCSF ChimeraX: Tools for structure building and analysis,” Protein Science, vol. 32, no. 11, p. e4792, 2023. _eprint: https://onlinelibrary.wiley.com/doi/pdf/10.1002/pro.4792.13

[42] “Case study: End-to-end processing of a ligand-bound GPCR (EMPIAR-10853) | CryoSPARC Guide.” https://guide.cryosparc.com/processing-data/tutorials-and-case-studies/case-study-end-to-end-processing-of-a-ligand-bound-gpcr-empiar-10853, Apr. 2025. 14

[43] D. Punjani, J. L. Rubinstein, D. J. Fleet, and M. A. Brubaker, “CryoSPARC: Algorithms for rapid unsupervised cryo-em structure determination,” Nature Methods, vol. 14, pp. 290–296, 2017. 17, 29

[44] D. Kimanius, B. Forsberg, S. Scheres, and E. Lindahl, “Accelerated cryo-em structure determination with parallelisation using GPUs in RELION-2,” eLife, vol. 5, e18722, 2016. 17

[45] T. Grant, A. Rohou, and N. Grigorieff, “cisTEM, user-friendly software for single-particle image processing,” eLife, vol. 7, p. e35383, 2018. 17

[46] T. Bepler, A. Morin, M. Rapp, J. Brasch, L. Shapiro, A. J. Noble, and B. Berger, “Positive-unlabeled convolutional neural networks for particle picking in cryo-electron micrographs,” Nature Methods, vol. 16, pp. 1153–1160, Nov. 2019. Publisher: Nature Publishing Group. 22, 27, 43

[47] T. Wagner, F. Merino, M. Stabrin, T. Moriya, C. Antoni, A. Apelbaum, P. Hagel, O. Sitsel, T. Raisch, D. Prumbaum, D. Quentin, D. Roderer, S. Tacke, B. Siebolds, E. Schubert, T. R. Shaikh, P. Lill, C. Gatsogiannis, and S. Raunser, “SPHIRE-crYOLO is a fast and accurate fully automated particle picker for cryo-EM,” Communications Biology, vol. 2, p. 218, June 2019. Publisher: Nature Publishing Group. 22

[48] F. Wang, H. Gong, G. Liu, M. Li, C. Yan, T. Xia, X. Li, and J. Zeng, “DeepPicker: A deep learning approach for fully automated particle picking in cryo-EM,” Journal of Structural Biology, vol. 195, pp. 325–336, Sept. 2016. 22

[49] J. Lehtinen, J. Munkberg, J. Hasselgren, S. Laine, T. Karras, M. Aittala, and T. Aila, “Noise2Noise: Learning image restoration without clean data,” Int. Conf. on Machine Learning, 2018. 24

[50] T. Bepler, K. Kelley, A. J. Noble, and B. Berger, “Topaz-Denoise: general deep denoising models for cryoEM and cryoET,” Nature Communications, vol. 11, p. 5208, Oct. 2020. Publisher: Nature Publishing Group. 24

[51] H. Li, H. Zhang, X. Wan, Z. Yang, C. Li, J. Li, R. Han, P. Zhu, and F. Zhang, “Noise-Transfer2Clean: denoising cryo-EM images based on noise modeling and transfer,” Bioinformatics, vol. 38, pp. 2022–2029, Mar. 2022. 24

[52] Q. Huang, Y. Zhou, H.-F. Liu, and A. Bartesaghi, “Joint micrograph denoising and protein localization in cryo-electron microscopy,” Biological Imaging, vol. 4, p. e4, Jan. 2024. 24

[53] A. Punjani, H. Zhang, and D. J. Fleet, “Non-Uniform Refinement: Adaptive regularization improves single particle cryo-em reconstruction,” Nature Methods, vol. 17, pp. 1214–1221, 2020. 31

[54] J. Zivanov, T. Nakane, and S. H. W. Scheres, “Estimation of high-order aberrations and anisotropic magnification from cryo-EM data sets in RELION-3.1,” IUCrJ, vol. 7, pp. 253–267, Mar. 2020. Publisher: International Union of Crystallography. 31

[55] J. Zivanov, T. Nakane, and S. H. W. Scheres, “A Bayesian approach to beam-induced motion correction in cryo-EM single-particle analysis,” IUCrJ, vol. 6, pp. 5–17, Jan. 2019. Publisher: International Union of Crystallography. 32

[56] Y. Z. Tan, P. R. Baldwin, J. H. Davis, J. R. Williamson, C. S. Potter, B. Carragher, and D. Lyumkis, “Addressing preferred specimen orientation in single-particle cryo-EM through tilting,” Nature Methods, vol. 14, pp. 793–796, Aug. 2017. Publisher: Nature Publishing Group. 32

